# Fast Generalized Longitudinal Functional Principal Components Analysis with Applications to Population-Level Neuronal Dynamics

**DOI:** 10.64898/2026.10.12.758639

**Authors:** Kelly A. Duffy, Nidhi Pai, David Schneck, Gleb Kozienko, Lvpiao Zheng, Annette Xenopoulos-Oddsson, Jing Wang, Luke A. Johnson, Jerrold L. Vitek, Mark Fiecas, Erjia Cui

**Affiliations:** Masonic Institute for the Developing Brain, University of Minnesota, Minnesota, USA; Division of Biostatistics and Health Data Science, University of Minnesota School of Public Health, Minnesota, USA; Department of Neurology, University of Minnesota School of Medicine, Minnesota, USA

**Keywords:** Neuronal Population Dynamics, Functional Data, Functional Principal Component Analysis, Deep Brain Stimulation

## Abstract

Functional data are widely collected across many fields. One approach for modeling functional data collected across repeated measures is longitudinal functional PCA (LFPCA), which allows for the decomposition of variability at multiple levels. However, existing estimation approaches for LFPCA can be computationally demanding for large or high-dimensional datasets. Here, we propose fast generalized LFPCA, a novel estimation technique that bins data along the functional domain and fits a marginal mixed model at each bin. Simulation results show that the proposed method is orders of magnitude faster than existing techniques while achieving similar accuracy, making LFPCA scalable to datasets with more densely sampled observations and larger sample sizes. We apply the proposed technique to model neuronal firing rate data both on and off deep brain stimulation (DBS) to investigate how the intervention modulates population-level neuronal activity. Our results suggest that the vast majority of between-neuron variation can be explained by four principal components, which relate to individual differences in neuronal firing as well as the arm manipulation paradigm under study. We also find that DBS significantly increases the variability of firing activity, likely driven by increased firing from a subset of neurons in response to DBS, a novel insight not obtained in previous analyses. Code to implement the proposed method and run simulation studies is included in the supplement.

## Introduction

### Background on Neuronal Firing Rate Analysis

Understanding the activity of individual neurons and how they work in concert within a large population of neurons is essential for elucidating the functional roles of neuronal populations and how they may be altered in many neurological disorders. It is also key for understanding the potential mechanisms underlying the efficacy of many available treatments, such as deep brain stimulation (DBS) (1). One way to investigate neuronal activity is by directly measuring the neuronal firing rate (FR) using single-cell recordings through microelectrodes across multiple neurons (2), which produces data from a population of neurons at a high-temporal resolution and has been widely used to understand brain structures and functional circuitry (3–5).

Many different analytical techniques have been proposed for analyzing FR data, with a large number of approaches relying on analyses of each neuron individually, such as by constructing simple tuning curves that measure neuronal firing rates across varying parameters of external stimuli (6–8). However, well-documented heterogeneity across single neurons can lead to difficulty in analysis and interpretation (9), often leading researchers to adopt simplifications such as analyzing only a subset of all recorded neurons or averaging across multiple trials. In addition, approaches that average across trials can obscure the true underlying neural time course when responses differ across trials, as is common in cognitive and motor tasks (10).

Although neuronal firing rate data provide a direct measure of neuronal activity and offer insights into its relationship with observed behavior, several characteristics of these data make analysis and interpretation particularly challenging. First, the FR data are generally not independent, single time point measurements, but rather are collected continuously and organized into temporal segments called “cycles” (or “trials”, when referring to a particular event or task) aligned to a predetermined reference time point. Given that FR data are *functional* in nature (11), an ideal analytical technique should appropriately account for this structure and the correlations it induces across time. Second, the FR data are generally not collected from a single neuron on a single occasion, but rather are collected repeatedly across time from many neurons, and are typically collected under different conditions. These study designs often result in a *multilevel* and longitudinal data structure. It is multilevel because we have multiple neurons (level-1) and multiple observations from a single neuron (level-2). The multiple observations over time also give the data a longitudinal structure, as the nature of the effect may change over the time course under study. Third, this multilevel, high-dimensional structure leads to an incredibly large amount of FR data. For example, data might be collected from a large number (e.g., hundreds) of neurons in thousands of repeated cycles within several data collection blocks or conditions that can in total comprise hours or days of FR information. Many existing analyses, which typically focus on a single neuron, fail to capture the high-dimensional, longitudinal, and multilevel structure of neuronal FR data. Indeed, very few analytical techniques can address all of these complexities, and it remains difficult to distill this information into a coherent, lower-dimensional form that can help answer scientific questions of interest, such as the effect of DBS interventions on neuronal activity in particular regions of the brain.

### Population Dynamics and Dimensionality Reduction

In recent years, there has been an increasingly strong interest in studying population-level neuronal activity (10, 12–15), which can reveal insights that cannot be gleaned from traditional single-neuron analysis, particularly when underlying mechanisms require coordination across multiple neurons. In addition to uncovering population-level features, such analyses can aid in hypothesis generation through exploratory data analysis (10). Population-level analytical approaches typically rely on some type of dimensionality reduction, assuming a shared latent space across multiple neurons. In doing so, these techniques can characterize how the firing of multiple neurons covaries coherently, while also separating out the noise that may occur from firing variability in the raw data. Principal Component Analysis (PCA) is by far the most common and general dimensionality reduction approach. PCA aims to identify a lower-dimensional space that captures most of the data variability, specifically by identifying a set of orthogonal directions through an eigendecomposition of the covariance matrix. However, directly applying PCA to raw FR data can be problematic because (1) it does not account for the functional nature of FR recordings over time; (2) it does not account for the correlations of recordings within a neuron. For this reason, it is common to average across trials (or cycles) or to temporally smooth neuronal firing data (10) before applying PCA. Rather than necessitating these transformations, which can themselves obscure effects of interest, we propose an innovative technique explicitly designed to accommodate data collected from multiple neurons and cycles: fast generalized longitudinal functional PCA (fast generalized LFPCA).

### Background on Longitudinal Functional Principal Component Analysis

Longitudinal Functional Principal Component Analysis (LFPCA) was proposed in Greven et al.(16) as an extension of the earlier Multilevel Functional Principal Component Analysis (MFPCA)(17). For the FR data, which longitudinally collects multiple cycles of functional data from multiple neurons, LFPCA is particularly well-suited over traditional PCA. Importantly, past work has demonstrated that when multilevel structure is present, performing a single-level FPCA can result in the identification of numerous additional spurious components due to the unaccounted for heterogeneity, decreasing parsimony and affecting the subsequent interpretation (17). Furthermore, the ability to separate out the variability at distinct levels of analysis is often of particular interest to researchers, as it provides insight into how changes differ at particular levels (e.g., between-neuron versus between-cycle) and can inform understanding of underlying mechanisms. While LFPCA has been used to model longitudinal functional data in several domains, including functional magnetic resonance imaging (fMRI) (18), accelerometer and activity (19), and diffusion tensor imaging (20), the original estimation technique is computationally expensive, especially for dense functional observations. Specifically, the covariance estimation process in (16) involves a separate linear regression for every pair of timepoints in the functional domain, followed by bivariate smoothing for each covariance function. In terms of the number of points on the functional domain *L*, the computational cost is on the order of *O*(*L*^2^), which can become prohibitive when *L* is large (i.e., with dense functional data).

To fill this critical gap, we propose an alternative estimation technique for the LFPCA model, namely fast generalized LFPCA. Instead of directly estimating the covariance functions, the proposed technique partitions the functional domain into bins and fits a marginal linear mixed model within each bin. This yields predictions of the neuron- and cycle-level random effects, which are used to calculate empirical covariance estimates on a smaller grid. The proposed procedure is orders of magnitude faster than the traditional technique with no substantial loss in recovery of simulated effects. For example, in one *n* = 300 simulation discussed below, the proposed method has a median runtime of roughly 2 seconds, compared to about 63 seconds for the existing method. This makes LFPCA feasible for longitudinal highdimensional data collected over thousands of visits and at thousands of time points per visit.

### Objective

In the present work, we will introduce the technical details of the proposed fast generalized LFPCA approach, and demonstrate how the approach can be used to address substantive questions of interest in neuroscience research. Specifically, we will apply the approach to the neuronal FR data to assess the effect of DBS on parkinsonian motor signs and neural activity in a non-human primate (NHP); see Section for data collection details. By applying fast generalized LFPCA, we will demonstrate how DBS affects population-level neuronal dynamics and alters the response to passive arm manipulation, which can potentially shed light on the underpinnings of sensorimotor deficits observed in Parkinson’s disease (PD), such as in sensory processing, as well how therapeutic DBS alters these dynamics. To our knowledge, this is the first study of the effect of DBS on population-level neuronal dynamics in parkinsonism.

The rest of the paper is organized as follows. In Section we describe our motivating dataset. The proposed fast generalized LFPCA model and implementation details are presented in Section . Section presents the simulation results. We present application results in Section and conclude with discussions in Section .

### Data

DBS in the subthalamic nucleus (STN) is an effective and common surgical treatment for PD(21–23). However, many questions remain regarding the effect of DBS on FR and neural activity in the basal ganglia-thalamocortical network in PD. This is particularly true for less commonly studied motor dysfunctions of PD, such as deficits in kinesthesia, or perception of limb movement, which may relate to deficits in sensorimotor processing. Here, we focus on the effects of STN DBS on the FR in the primary motor cortex (M1) that coincides with a passive arm manipulation paradigm. Throughout this section, we will reference panels of Figure 1, which provides an overview of our data structure.

**Fig. 1.**
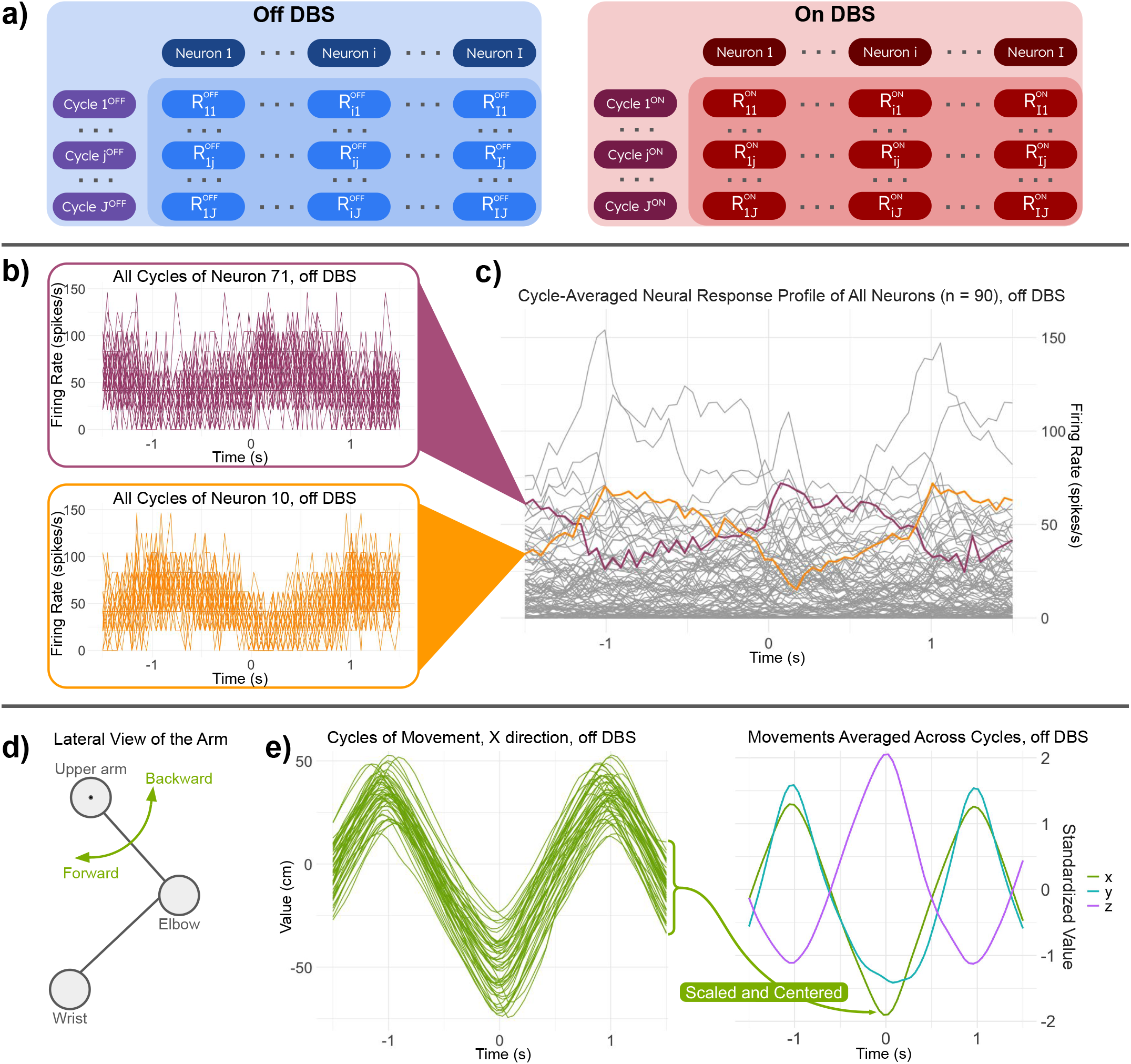
The overall data collection paradigm and data structure. **(a)** The data structure within the two condition blocks of data collection: prior to DBS (“OFF DBS”) and during DBS (“ON DBS”). Data are collected from multiple neurons and across multiple cycles. **(b)** The firing rates of two representative neurons across all cycles in the OFF DBS condition. **(c)** The firing rate, averaged across all cycles in the OFF DBS condition, for neurons 71 (purple) and 10 (orange), and the overall average firing rate of the other 88 neurons in gray. **(d)** The passive arm manipulation paradigm that was conducted simultaneously with the neuronal firing data collection. The arm of the NHP was moved forwards and backwards from the shoulder, and arm position information was collected via a sensor on the wrist. **(e)** Arm position information collected in the paradigm depicted in (d). The left panel displays the arm position, in the X (anterior-posterior) direction across all cycles in the OFF DBS condition. The right panel displays the average of the X direction position across all cycles in the condition block, along with the average Y (superior-inferior) direction position, Z (lateral-medial) direction position across the cycles.

#### Data Collection

We begin by outlining the data collection approach in our motivating neuronal FR data. One NHP (macaca mulatta, 13 y.o.) was rendered parkinsonian by administrating MPTP (1-methyl-4-phenyl-1,2,3,6-tetrahydropyridine). A DBS lead (NuMed) was implanted in the right STN. Neural signals were collected both prior to DBS initiation and on STN DBS (130Hz, 0.2 mA) from the arm area of primary motor cortex (M1), using a 96-channel Utah microelectrode array (Blackrock Microsystems) and a TDT workstation (Tucker Davis Technologies, ∼24 kHz sampling rate). Details of these procedures are described in (24). Neuronal data were then band pass filtered at 300-5000 Hz and sorted in Offline Sorter software to extract single unit spike firing data (a total of 90 spike trains). The neuronal data were collected under a passive manipulation paradigm of left shoulder flexion and extension. Arm kinematics were monitored using a motion capture system (Motion Analysis, 100 Hz frame rate) with a reflective marker placed over the left wrist. Kinematic data were then processed in Matlab to extract wrist location and movement.

Data from spike firing and movement recordings that were collected prior to STN DBS are grouped together and referred to as the “OFF DBS” condition. Data collected during STN DBS are also grouped together and referred to as the “ON DBS” condition. Continuous measurements of both spike firing and movement data were split into 3-second intervals which we call “cycles”. Movement cycles were aligned at the flexion-extension transition point (that is, the time of the maximum shoulder extension movement was detected and defined as 0) such that there were 1.5 seconds of data before this point and 1.5 seconds after this point. Data were then discretized into 50 millisecond (ms) increments by downsampling the kinematic data to 20 Hz and binning the neuronal data into 50 ms intervals to calculate the neuronal FR at the same frequency.

For the FR data, *I* = 90 neurons were considered. For the OFF DBS condition, there were *J*^*OF F*^ = 55 cycles of data collected, and for the ON DBS condition there were *J*^*ON*^ = 32 cycles. Data were collected from the same neurons in both conditions. Thus, within each condition block, we observe firing rates *R*_*ij*_ for neuron *i* and cycle *j*. This structure is illustrated in panel a) of Figure 1. To further clarify our data structure, we present panels b) and c) of Figure 1. Panel b) shows 55 cycle-specific FR recordings in the OFF DBS condition for 2 randomly selected neurons. For the subpanels presenting the data of each of those neurons, the *x*-axis represents time (that is, the 3-second interval that comprises the cycles described above), and the *y*-axis represents the firing rate in spikes per second. Given that there are multiple cycles of FR data collected for each neuron, there is within-neuron, between-cycle variability present in the data. Panel c) displays the average FR trajectories across the cycles in the OFF DBS condition for the two selected neurons, as well as for other 88 neurons under study, with axes having a same interpretation as in panel b). This demonstrates the between-neuron variability within each data collection condition.

For the kinematic data, the passive manipulation paradigm is illustrated in panel d) of Figure 1. From this paradigm, arm kinematic data was tracked using the motion capture system previously described to produce arm location data, which is illustrated for the “X” direction of movement (roughly, the anterior-posterior direction). Kinematics were also tracked in the “Z” (approximately laterial-medial) and “Y” (superior-inferior) directions. The arm position in each of these directions can be averaged across the cycles within each data condition to form a condition-specific average arm trajectory, as visualized in panel e) of Figure 1 for the OFF DBS condition. In summary, we have data that are functional in nature, as the FR is collected continuously over time at each cycle. Furthermore, because multiple cycles of data were collected within a neuron in each condition block, our data has a clear longitudinal structure. This makes our data a perfect candidate for the proposed fast generalized LFPCA approach, which allows us to decompose the variation occurring between individual neurons as well as between cycles within a neuron. In this way, fast generalized LFPCA enables us to investigate population-level neuronal dynamics, and comparisons between the ON DBS and OFF DBS conditions allow us to explore the effect of DBS on these population-level dynamics.

#### Data Preprocessing

Prior to fitting the LFPCA model, a log(*R*_*ij*_(*t*) + 1) transformation was applied to the raw neuronal FR data. There is evidence that some neurons exhibit log-normally distributed firing rates (25, 26), and the log transformation has been widely used in the statistical analysis of firing rates (27–29). For this project, log transformation was selected for its relative simplicity and the less skewed resulting distribution of residuals.

## Methods

### Longitudinal Functional Principal Component Analysis (LFPCA) Model

We first outline the LFPCA model (16). Consider longitudinal functional data of the form *R*_*ij*_(*s*) for neuron *i* = 1, …,*I* and cycle *j* = 1, …, *J*_*i*_ at time *s* ∈ *{s*_1_,…, *s*_*L*_*}* ∈ *S*, where *S* is a compact domain. Note that each cycle has *L* observations at the same set of time points. In our application, each neuron has the same number of cycles *J*_*i*_ = *J* within each condition, but this is not required. Cycle *j* is observed at time *T*_*ij*_. Consider the longitudinal functional model 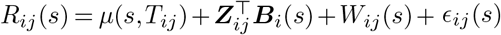, where *µ*(*s, T*_*ij*_) is the population mean function, ***B***_*i*_(*s*)= (*B*_*i*,0_(*s*), *B*_*i*,1_(*s*))^*T*^ is the vector of neuron-specific (in other contexts, subject-specific) functional random effects with corresponding design matrix ***Z***_*ij*_ = (1, *T*_*ij*_)^*T*^, *W*_*ij*_(*s*) is the neuron-cycle (in other contexts, subject-visit) functional random effect, and *E*_*ij*_(*s*) is random white-noise error with a variance of *σ*^2^. Here, *B*_*i*,0_(*s*) and *B*_*i*,1_(*s*) are the neuron-specific functional intercept and slope, respectively. The random functions ***B***_*i*_(*s*) and *W*_*ij*_(*s*) are assumed to be mutually independent mean-0 random processes. Let *K*_0_(*s, u*) = cov *{B*_*i*,0_(*s*), *B*_*i*,0_(*u*) *}* and *K*_1_(*s, u*) = cov *{B*_*i*,1_(*s*), *B*_*i*,1_(*u*) *}* be the covariance functions of *B*_*i*,0_(*s*) and *B*_*i*,1_(*s*), respectively, and let *K*_01_(*s, u*) = cov {*B*_*i*,0_(*s*), *B*_*i*,1_(*u*) } be their cross-covariance function. *K*_0_(*s, u*) and *K*_1_(*s, u*) are symmetric while *K*_01_(*s, u*) is not. Then 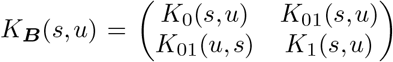 and *K*_*W*_ (*s, u*)= cov *{W*_*ij*_(*s*), *W*_*ij*_(*u*)} give the between-neuron and within-neuron variance, respectively.

Following Mercer’s theorem (30), the between-neuron covariance function *K*_***B***_(*s, u*) can be eigendecomposed as 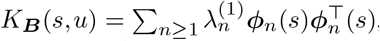, where 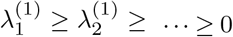 are the level-1 eigenvalues associated with the eigenfunctions 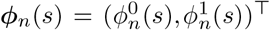. The set of eigenfunctions *{****ϕ***_*n*_(*s*)*}* forms an orthonormal basis with respect to the additive scalar product 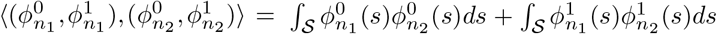. Following the Karhunen-Loeve theorem (31), the neuron-specific random functions ***B***_*i*_(*s*) can be written as ***B***_*i*_(*s*)= ∑ _*n*≥ 1_ *ξ*_*in*_***ϕ***_*n*_(*s*), where *ξ*_*in*_ are mutually uncorrelated scores with a mean of zero and a variance of 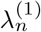. The within-neuron (between-cycle) variability can be eigendecomposed in a similar fashion as 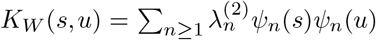 where 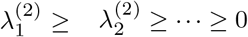are the level-2 eigenvalues that have associated orthonormal eigenfunctions *ψ*_*n*_(*s*). The neuron-cycle-specific random functions *W*_*ij*_(*s*) can then be written as *W*_*ij*_(*s*)= ∑_*n* ≥1_ *ζ*_*ijn*_*ψ*_*n*_(*s*), where *ζ*_*ijn*_ are mutually uncorrelated scores with a mean of zero and a variance of 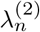. To achieve dimensionality reduction, the first several principal components are kept at each level, where the number can be chosen, for example, based on the proportion of variance explained (PVE) (17, 32). Through such decomposition, LFPCA reduces the longitudinal functional data into two sets of uncorrelated scores: the level-1 (between-neuron) scores 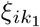 and the level-2 (within-neuron, between-cycle) scores 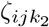. We also obtain eigenfunctions at level-1 and level-2, which show patterns of variation in the functional data at each level.

LFPCA is well-suited to model the repeated measures from multiple neurons in our motivating FR data, as it allows the explicit investigation of population-level neuronal activity (by focusing on the level-1 eigenfunctions) while appropriately accounting for the heterogeneity between individual neurons and across cycles within neurons. However, the application of the approach has been limited in the past due to computational constraints. Specifically, the original LFPCA approach (16) requires estimating each covariance function on the full *L* by *L* grid of time points, followed by bivariate smoothing. As *L* increases, computation time increases significantly. Here, we propose an alternative procedure for fitting the LFPCA model, which is much faster for large *L*. As described in Section, instead of directly estimating and smoothing covariance functions, the data is binned along the functional domain, and a linear mixed model is fit at each bin (33). Because the variance components and predictions admit closed-form estimates and the empirical covariances are estimated and smoothed on a reduced grid of *Q* bin centers, the computational cost scales with the number of bins *Q* and leads to substantial computational savings, as shown in Section .

#### Fast Generalized LFPCA

In this section, we describe the fast generalized LFPCA, whose steps are outlined in Algorithm 1. We begin by estimating and removing the population mean function (Section ). Next, we partition the functional domain into *Q* overlapping bins and fit a linear mixed model (LMM) within each bin to obtain predictions of the neuron-and cycle-level random effects (Section ). From these predictions, empirical covariance estimates are calculated on the reduced *Q × Q* grid of bin centers; these estimates are smoothed and decomposed to obtain eigenvalues and eigenfunctions of the covariance functions (Section ). After obtaining the mean function, eigenvalues, and eigenfunctions, the model reduces to a mixed effects model and we predict scores using Henderson’s mixed model equations (MME; Section ).

##### Algorithm 1

Fast Generalized LFPCA Algorithm

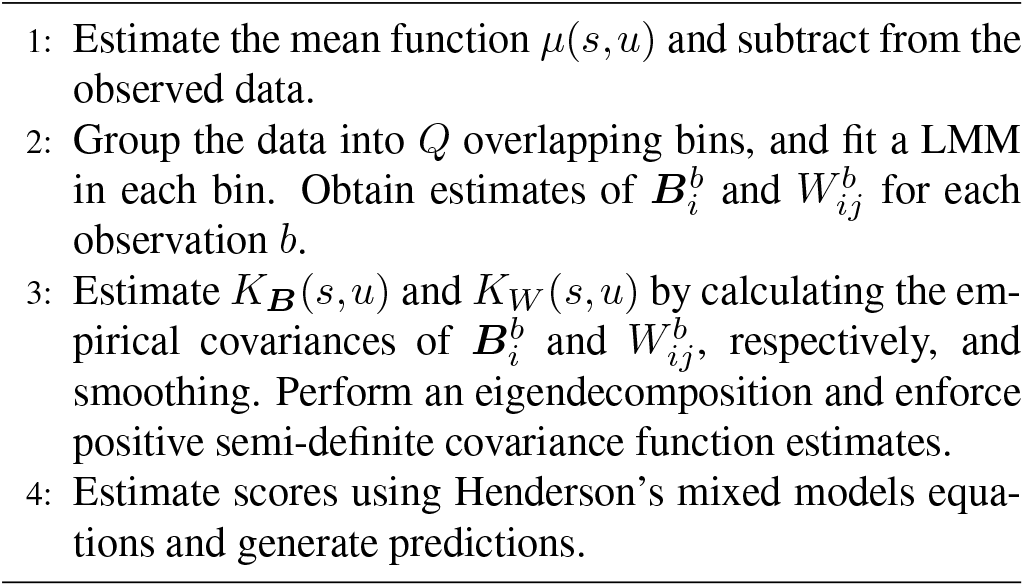

##### Fixed effects estimation

We first obtain an estimate of *µ*(*s, T*_*ij*_) under a working independence assumption using a tensor product of P-splines, although different fixed effect structures and estimation frameworks can be used based on the application. After obtaining an estimate 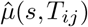, we subtract the estimate from the functional data and denote the residuals as 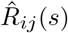.

##### Marginal linear mixed models

Fast generalized LFPCA diverges from existing LFPCA methods in the covariance estimation process. The data is first grouped into *Q* overlapping bins, each of width *w*, with centers evenly spaced on *S*. A bin that extends past the edge of *S* is truncated, so its width is smaller than *w*. The parameters *Q* and *w* are calibrated based on the application. For *b* = 1, …, *Q*, let *L*_*b*_ denote the number of grid points among *s*_1_, …, *s*_*L*_ that fall in bin *b*. We further let *l* index the grid points within a bin, so that 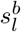 denotes the *l*th grid point in bin *b*, for *l* = 1, …, *L*_*b*_.

After binning, a linear mixed-effects model (LMM) is fit in each of the *Q* bins. Specifically, the model for bin *b* has the form 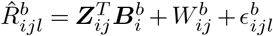, where 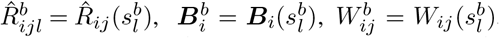, and 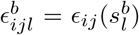 restrict their respective quantities to the points in bin *b*. Note that an observation can belong to multiple bins since the bins overlap. The working assumptions of the model are 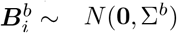 where 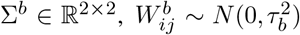, and 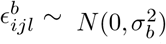. The neuron-level, cycle-level, and error terms are assumed to be mutually independent, and independent over neurons and cycles.

This model can be easily implemented using existing LMM software, such as `lme4`(34) in `R`. However, only a few quantities are needed from the model fit, which can be obtained even more quickly using closed-form estimates; details are in Supplementary Section S1. We use ANOVA-style estimators for the variance components 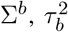, and 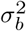. Henderson’s mixed models equations (MME) are used for predicting 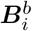 and 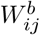 in the marginal LMMs.

##### Covariance estimation and eigendecomposition

Using the predictions 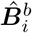 and 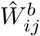 from each bin, we now construct estimates of *K*_***B***_(*s, u*) and *K*_*W*_ (*s, u*). Define 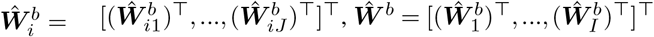, and 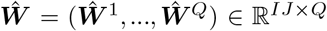; define 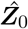 and 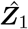 analogously. The four covariance functions are estimated by the sample covariances 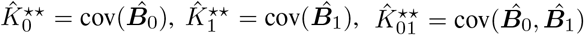, and 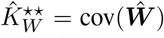, each evaluated on the grid of bin centers rather than on *s*_1_, …, *s*_*L*_. A bivariate smoother is then applied to the four estimates, yielding 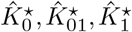, and 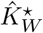 on the full grid *s*_1_, …, *s*_*L*_. Any bivariate smoother can be used here; we use a tensor product of P-splines with 16 basis functions per dimension in the application.

After smoothing, the estimates are eigendecomposed to obtain positive semi-definite (PSD) covariance functions and enable KKL expansions. For the neuron-level covariance function *K*_***B***_(*s, u*), the estimated smoothed covariance functions are first concatenated into a block matrix 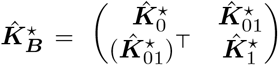 whose eigendecomposition gives eigenvalues 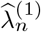 and eigenvectors 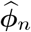 for *n* = 1, …, *L*. Following the literature (16, 32), we trim negative eigenvalues to obtain the PSD estimate 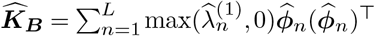. Estimates of *K*_0_(*s, u*), *K*_01_(*s, u*), and *K*_1_(*s, u*) are the corresponding blocks of 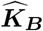. The within-neuron, across cycle-level proceeds identically: eigendecomposing 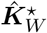 gives eigenvalues 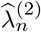 and eigenvectors 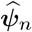 for *n* = 1, …, *L*, and trimming yields 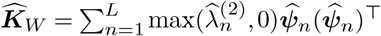.

Finally, the dimensionality reduction is achieved by truncating each expansion based on PVE, retaining the first *N*_*B*_ neuron-level principal components; *N*_*B*_ is chosen large enough such that 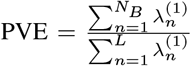 exceeds prespecified threshold. Each vector 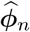 contains two blocks of *L* entries each. Estimates for 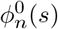 are given by entries 1 to *L*, while estimates for 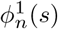 are given by entries *L* +1 to 2 *L*. Similarly, we retain *N*_*W*_ cycle-level components based on PVE.

##### Scores and prediction

After estimating the mean function, eigenfunctions, eigenvalues, and error variance, the longitudinal functional model in Section reduces to a mixed effects model, and we predict the scores *ξ*_*in*_ and *ζ*_*ijn*_ using MMEs. Substituting the estimated mean function, eigenfunctions, and scores back into the model yields predictions of *R*_*ij*_(*s*) at any point in *S*.

#### Generalized outcomes

Because the estimation is built on linear mixed models, the model easily extends to non-Gaussian outcomes via generalized linear mixed models (GLMM) and the associated fitting tools. Assume *R*_*ij*_(*s*) ∼ EF {*m*_*ij*_(*s*)}, where EF denotes an exponential family distribution. The conditional mean *m*_*ij*_(*s*) is modeled through a link function *g*( *·*) as 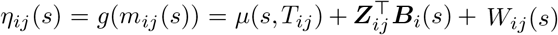. As with Gaussian outcomes, the data is binned into *Q* overlapping bins, and the GLMM for bin *b* takes the form 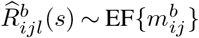 with 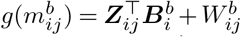. Although the closed-form estimators in Section for the variance components and predictions no longer apply, the marginal models can be fit easily using standard GLMM algorithms such as penalized quasi-likelihood, Laplace approximation, or adaptive Gauss-Hermite quadrature (35). These algorithms are available in the `R`functions `glmmPQL`in `MASS`(36), or `glmer`in `lme4`(34)

### Simulations

#### Simulation design

We conduct extensive simulations to evaluate the performance of the proposed fast generalized LFPCA in estimating eigenfunctions and covariance functions, benchmarking it against the original LFPCA method (16), hereafter referred to as “Greven et al.’s approach”. Data is generated according to the Karhunen-Loeve decomposition 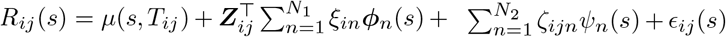. We set *I* = 90 neurons (in other contexts, subjects), *J* = 48 cycles per neuron (in other contexts, visits per subject), and *L* = 62 points on the functional domain (time points per cycle). The parameters are set to mirror the observed FR data. Since the focus is on random effects, we set *µ*(*s, T*_*ij*_) = 0 for simplicity. The *N*_1_ = 2 eigenfunctions for the neuron-level functional intercept are 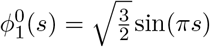 and 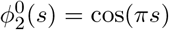, and the neuron-level slope eigenfunctions are 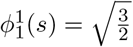 and 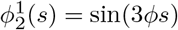. The cycle-level functions are *ψ*_1_(*s*) = 1 and 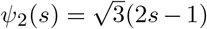. The neuron-level eigenvalues are set to 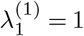 and 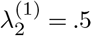, and the cycle-level eigenvalues are 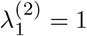 and 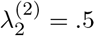. The cycle times *T*_*ij*_ are generated from a Unif[0, 1] distribution. The scores are generated from a centered normal distribution with variance equal to the eigenvalues: 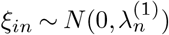 and 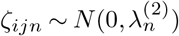. Finally, the error *E*_*ij*_(*s*) is generated from a *N* (0, *σ*^2^) distribution where *σ*^2^ = 1. For each simulation scenario, 300 datasets are simulated.

When fitting the models, we vary the number of bins *Q* = 5, 10, 25, 50 and the bin width *w* = .05, .1, .15,.2,.3,.4. The result is omitted if the combination of bin width and number of bins does not cover the functional domain. We also include a version of the model without binning; that is, each point *s*_*l*_ is in its own bin. In this case, the cycle-level effects are not identifiable. The fit is evaluated using the integrated squared error (ISE) of the eigenfunctions, ISE of the covariance functions, and the running time. All models are fit in `R`, and the code to run the simulations accompanies this paper.

#### Simulation results

The distribution of ISE of the estimated eigenfunctions in simulations is shown in Figure 2, where the first, second, and third columns correspond to the neuron-level intercept, the neuron-level slope, and the cycle-level intercept. The rows correspond to the two eigenfunctions estimated at each level. For the neuron-level eigenfunctions, the proposed fast generalized LFPCA method, with any bin width and number of bins, has a lower median ISE than Greven et al.’s approach. For both of the first neuron-level eigenfunctions (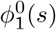 and 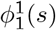), all variants of the proposed method perform similarly. For the second neuron-level intercept eigenfunction 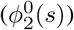, using 10 bins has a slight advantage over the increasing number of bins. For the second neuron-level slope eigenfunction 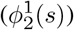, the smaller bin widths *w* = .05 and *w* = .1 have lower median ISE than larger bin widths. For the within-neuron eigenfunctions, Greven et al.’s approach outperforms the proposed method in terms of median ISE. However, the results are comparable when using a larger bin width (*w* = .2) and *Q* = 10 bins. This is expected as a larger bin width allows more observations from the same neuron and cycle to be included in each marginal LMM. Supplementary Figure S1 shows the ISE of estimated covariance functions in simulation. Overall, the patterns are similar to that of the eigenfunctions in Figure 2.

**Fig. 2.**
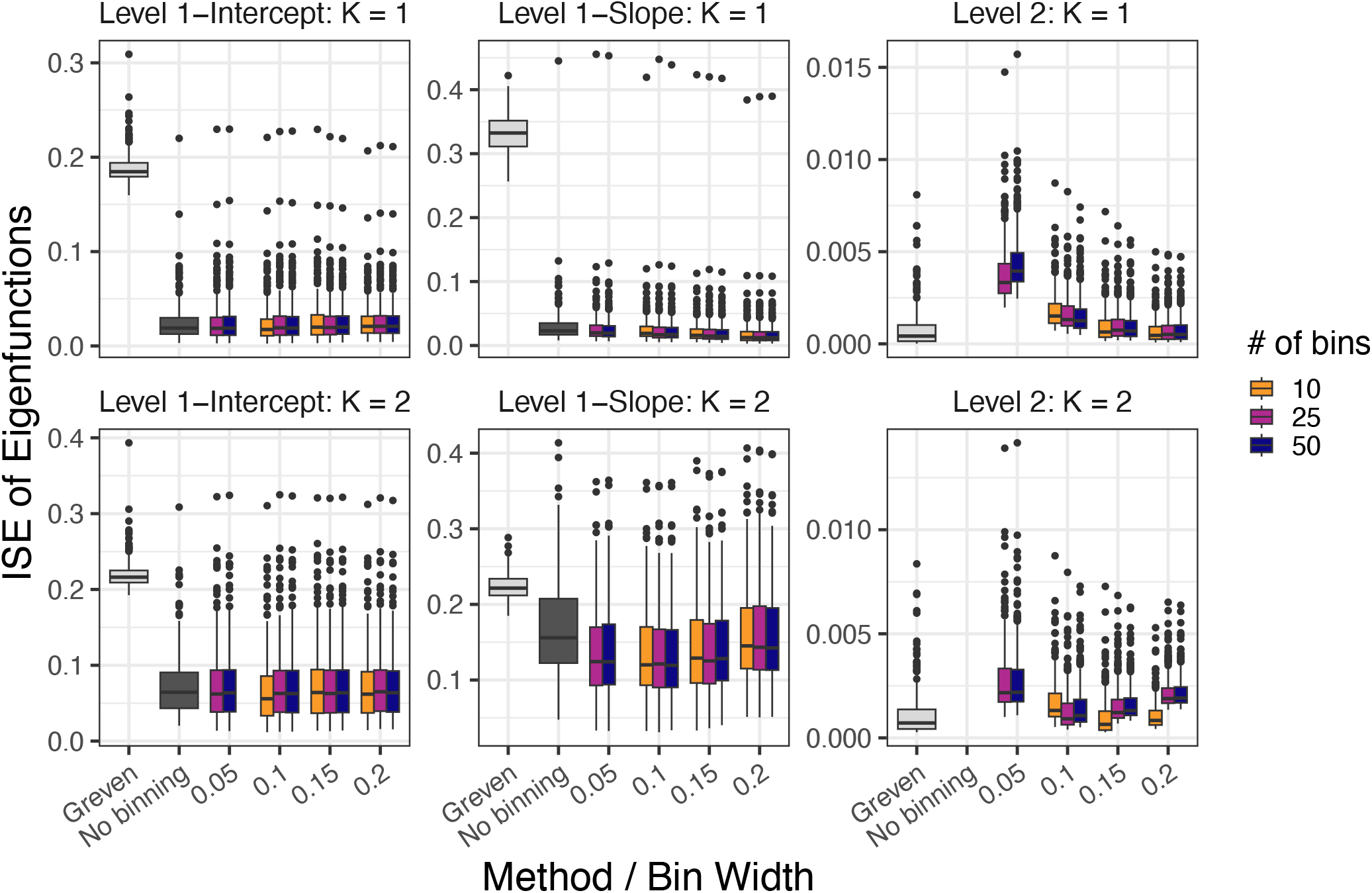
Boxplots of integrated squared error (ISE) for eigenfunctions in simulation. From left to right, the columns correspond to the neuron-level intercept, neuron-level slope, and cycle-level intercept, and the rows correspond to the top two eigenfunctions.

Figure 3 shows the runtime of the fast generalized LFPCA and Greven et al.’s method. Notably, all variants of our proposed method run significantly faster than Greven et al.’s method. For example, with 10 bins and a bin width of .15, the median runtime of fast generalized LFPCA for *n* = 300 is 1.97 seconds, compared to 62.68 seconds for Greven et al.’s method. A higher number of bins results in a slower runtime; with 50 bins and a width of .15, the median runtime is 4.97 seconds. However, this is still more than 10 times faster than Greven et al.’s method.

**Fig. 3.**
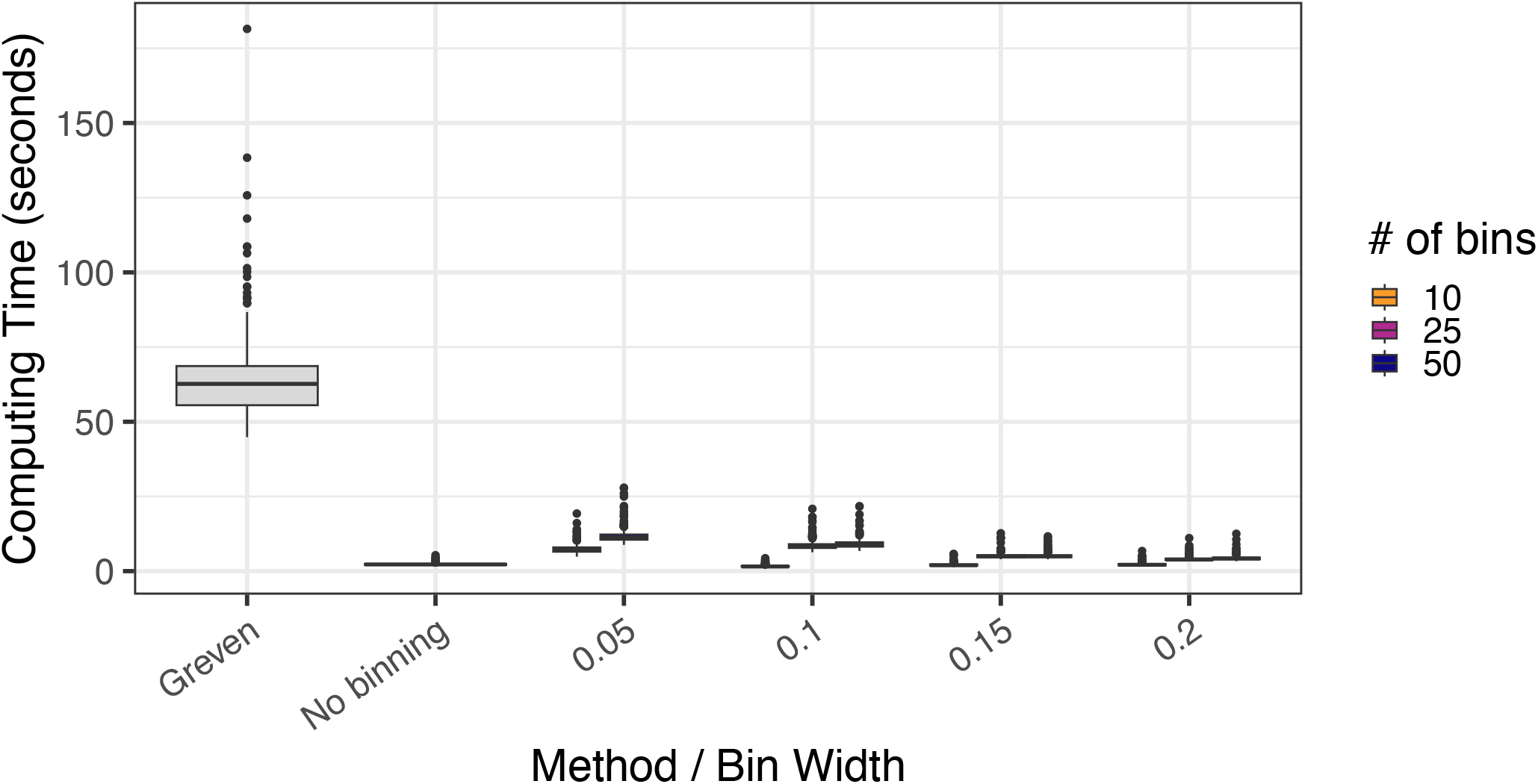
Boxplots of the runtime of Greven et al.’s(16) method and variants of the proposed method.

In summary, our simulations show that the proposed method is substantially faster than the existing method with no substantial loss in performance. This makes our fast generalized LFPCA particularly appealing for large-scale datasets, which are becoming increasingly common in neuroscience and other fields.

### Application to Neuronal Firing Rate Data

#### Fitting Fast Generalized LFPCA to FR Data

In our application, we fit separate fast generalized LFPCA models for the two data conditions (ON and OFF DBS), in which the neurons were treated as level 1 units and the multiple cycles of FR were treated as level 2 units. In other words, level 1 here represents the “between-neuron” variability, which consists of both a level 1 intercept and a level 1 slope over the course of each cycle, and level 2 represents the within-neuron, “between-cycle” variability. We select the number of eigenfunctions to explain at least 99% of the total variability at both level 1 and level 2. Based on the above simulation results, we elected to fit the LFPCA models using *Q* = 25 bins and a bin width *w* = 0.20.

To evaluate each LFPCA model, we illustrate and examine the first four eigenfunctions, which explain the greatest amount of variance at both level 1 (between-neuron) and level 2 (between-cycle) within each condition. Note that while the PVE is estimated jointly for the two between-neuron effects (intercept and slope), the eigenfunctions are estimated and plotted separately. To evaluate whether STN DBS has an impact on the variance across the on DBS and off DBS conditions, we also report the total variance across the two conditions, as well as the relative proportion of the overall variance explained at level 1 and level 2.

#### Few Eigenfunctions Explain Most Neuronal Variability

Table 1 provides the within-level PVE of the first four eigenfunctions for level 1 and level 2 for the two stimulation conditions (OFF DBS and ON DBS). Figures 4 and 5 display those first four eigenfunctions at level 1 (both intercept and slope). Here, we will focus on the results at level 1, which represent the between-neuron effects that are our primary interest. The first four eigenfunctions at level 2 are presented in Supplementary Material (Figures S2 and S3).

**Table 1.** Within-level percentage of variance explained (PVE) per principal component for the OFF and ON DBS conditions. Each component’s PVE is normalized by the total variance within its own level. Level 1 denotes the between-neuron level, and Level 2 denotes the within-neuron (between-cycle) level.

| Block | Level | PC1 | PC2 | PC3 | PC4 |
| --- | --- | --- | --- | --- | --- |
| Off DBS | Level 1 | 88.7% | 5.7% | 3.3% | 0.7% |
|  | Level 2 | 18.2% | 11.5% | 11.4% | 9.7% |
| On DBS | Level 1 | 85.9% | 4.7% | 4.3% | 2.6% |
|  | Level 2 | 26.6% | 14.3% | 9.1% | 8.2% |

**Fig. 4.**
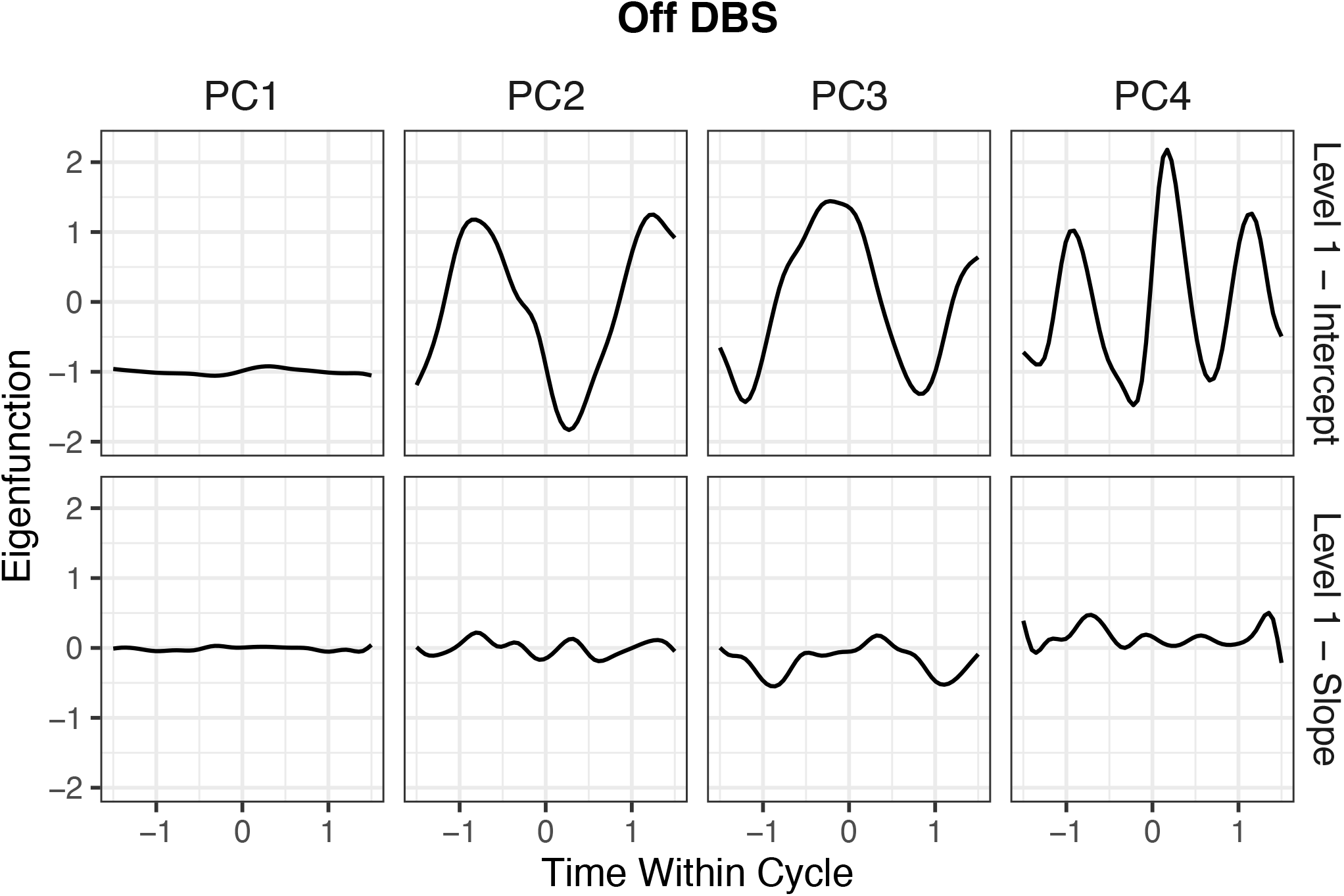
The first four eigenfunctions (PCs) explaining the largest proportion of variance at level-1 of the OFF DBS condition of activity. The top row displays the first four intercept eigenfunctions at level-1 (the between-neuron intercepts), and the second row displays the first four slope eigenfunctions at level 1 (the between-neuron slopes).

**Fig. 5.**
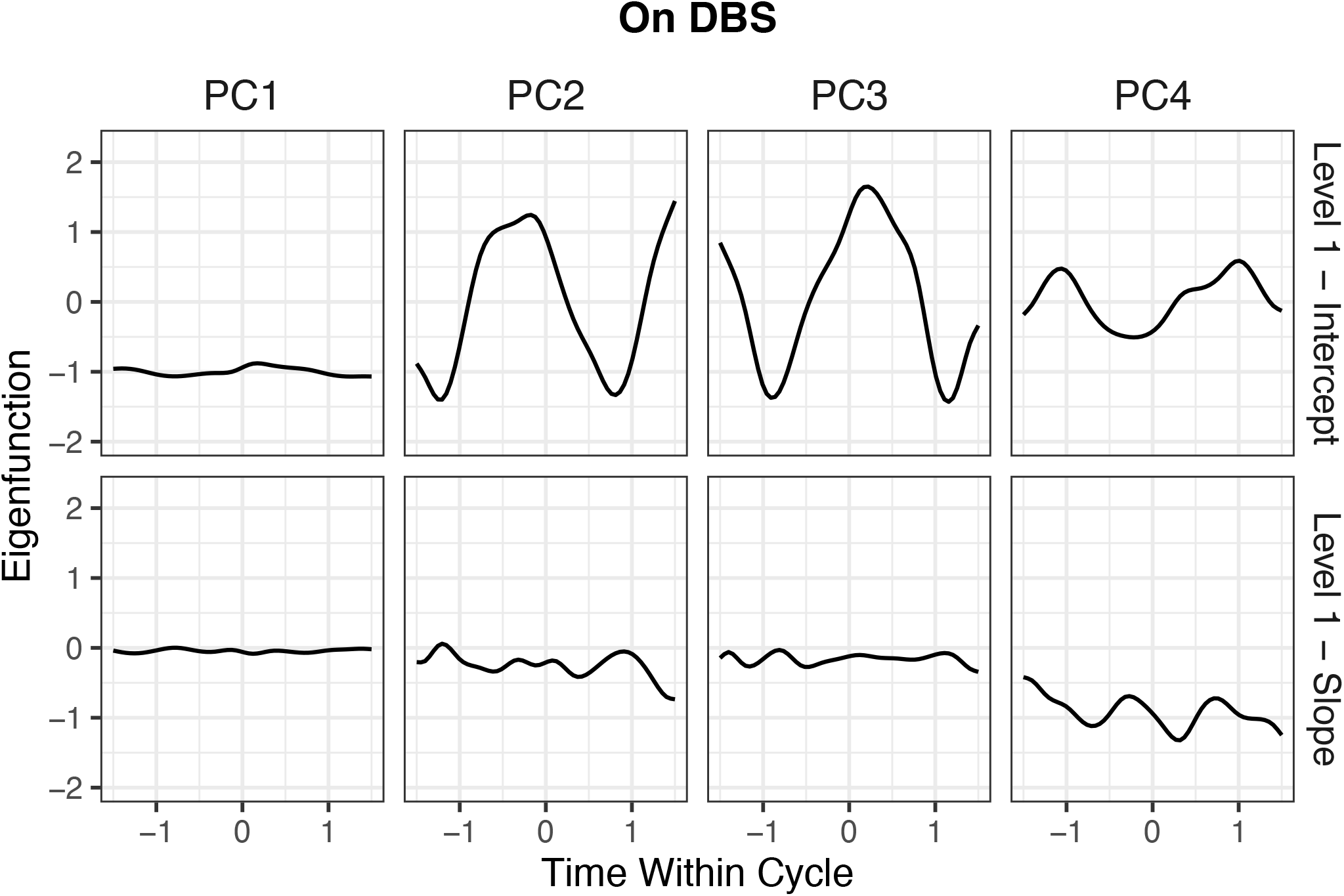
The first four eigenfunctions (PCs) explaining the largest proportion of variance at level-1 of the OFF DBS condition of activity. The top row displays the first four intercept eigenfunctions at level-1 (the between-neuron intercepts), and the second row displays the first four slope eigenfunctions at level-1 (the between-neuron slopes).

For both conditions, the first eigenfunction (“PC1”) explains approximately 85%-90% of the total variance at level 1. For the level 1 intercept, the first eigenfunction exhibits a relatively flat shape across the time span for both the ON and OFF DBS conditions. This eigenfunction can be interpreted as an overall indicator of FR differences between neurons. Indeed, there was great variability in which neurons fired in response to the passive arm manipulation, both in terms of absolute firing and the degree of the response, and thus the greatest amount of variability between neurons is explained by overall neuronal firing in response to passive arm manipulation. In effect, the first eigenfunction is operating similarly to a neuron-specific, non-functional random intercept. For both conditions, the next two eigenfunctions (“PC2” and “PC3”) explain a much smaller proportion of variance, approximately 1%-6% of the total. In both conditions, these two components have a very similar sinusoidal shape, with a similar period and frequency but with a phase shift between them due to the orthogonality constraint. Given the shape, we hypothesize that PC2 and PC3 relate to FR behavior under the passive arm manipulation paradigm. PC4 explains less than 1% of the total in variance in level-1 for the OFF DBS condition, and about 2.5% of the variance in the ON DBS condition. The shape of the eigenfunction differs between the two conditions, which, combined with the relative difference in PVE, suggests that PC4 may be more substantively meaningful in the ON DBS condition. We further explore the potential relationship between the PCs and the passive arm manipulation paradigm by investigating the correlations between the estimated eigenfunctions and the kinematic arm position information in the Supplementary Materials (Section S4.2). The results overall suggest that DBS does modulate the relationship between neuronal firing and arm position, by modifying both whether neuronal firing relates to arm position at all and increasing selectivity in which facets of arm motion relate to firing activity.

For the level-1 slope eigenfunctions, which represent the neuron-specific slope over the time course, all four first components in both conditions are relatively flat across the cycle and centered around zero. This suggests that there is not strong evidence of a neuron-specific longitudinal effect on neuronal firing across the cycles within either the ON or OFF DBS condition. There is a small decline in PC4 in the ON DBS condition, suggesting that some neurons may show a slight decrease (or increase) in firing over the course of the repeated cycles, but this component explains a relatively small proportion of the variance (2.6% at level-1).

The eigenfunctions for the level-2 components are presented in Supplementary Figures S2 and S3. The first eigenfunction explains a much smaller relative proportion of variance at level 2 than at level 1, with PVE values of 25%-30%. As a result, a much larger number of eigenfunctions was required to explain a majority of the total variance at level 2: 16 components (for both the OFF and ON DBS conditions) were needed to explain 99% of the variance at level 2, versus 5 components (for the OFF DBS condition) and 7 components (for the ON DBS condition) at level 1. “PC1” at level 2 has a somewhat similar overall shape to “PC1” at level 1: a fairly flat shape across time that effectively acts as an offset from the mean. Given that this is a within-neuron, between-cycle effect, we interpret this component as difference in overall FR across particular cycles of data collection within the neurons. PCs 2 through 4 are much more dissimilar across the two conditions at level 2 than the PCs in level 1. For the OFF DBS condition, they are generally highly nonlinear functions that may explain neuron- and cycle-specific deviations in firing patterns. For the ON DBS condition, we see one function showing a sustained increase across the data collection cycle (“PC2”) as well as two sinusoidal functions (“PC3” and “PC4”) that are somewhat similar to the intercept functions at level 1, which likely relate to the passive arm manipulation. This suggests that during DBS there may be variation across cycles in how some neurons fire in response to the passive arm manipulation paradigm.

#### Variance Increases During DBS

We additionally explore the total variation attributed to each of the two levels (level 1, the between-neuron effects, and level 2, the within-neuron, between-cycle effects) in the LFPCA models across the OFF DBS and ON DBS conditions. Results are shown in Table 2. We observe a nearly 25% increase of the total variance from the OFF DBS condition (1.98) to the ON DBS condition (2.45). Looking at the relative contribution of level 1 and level 2, the proportion explained by level 2 is much higher on DBS (17.55%) than off DBS (8.08%). These results suggest that DBS substantially increases the overall variance of the neuronal FR, with variability increasing at both levels but most prominently at the within-neuron, between-cycle level. We want to highlight that the separation of the variability at level 1 (between-neuron) and level 2 (within-neuron) is a distinct advantage of the LFPCA approach. In a standard PCA or functional PCA (FPCA), these two distinct sources of variability would be estimated together, and this key insight into how variability and the firing patterns of neurons is changing in response to DBS would be lost.

**Table 2.** Variance at each level (level 1, between-neuron, and level 2, within-neuron across cycles) and overall in the OFF DBS and ON DBS conditions. For level 1 (first column) and level 2 (second column) the overall variance is given, followed by the relative proportion of variance at that level (relative to the total variance, given in the third column) in parentheses.

|  | Level 1 Variance (Proportion) | Level 2 Variance (Proportion) | Total Variance |
| --- | --- | --- | --- |
| Off DBS | 1.82 (91.92%) | 0.16 (8.08%) | 1.98 |
| On DBS | 2.02 (82.45%) | 0.43 (17.55%) | 2.45 |

#### Single-Neuron Analysis Supports LFPCA Findings

To better understand and interpret the findings of how variability changes, both overall and at levels 1 and 2, we conducted a post-hoc analysis to assess variability at the single-neuron level. Using the firing data from *I* = 90 individual neurons, we calculate across-neuron variability (measured as the standard deviation of the FR across neurons at each time point within each cycle, averaged across cycles) and across-cycle variability (measured as the standard deviation of the FR across all cycles within each neuron). The effect of the DBS condition on variability is evaluated by conducting Wilcoxon rank-sum and signed rank tests (37). We also evaluate, at the individual-neuron and individual-cycle level, how this variability correlates with the mean FR, for both across-neuron variability (where the mean FR across all neurons is calculated for each cycle) and across-cycle variability (where the mean FR across all cycles is calculated for each neuron). We additionally compare the within-neuron mean FR (taken across cycles and averaged over time) and variability (specifically, the standard deviation across cycles, averaged over time) at the individual-neuron level to more directly compare activity ON and OFF DBS.

In evaluating across-neuron variability, a Wilcoxon rank-sum test confirmed that the ON DBS condition had significantly higher across-neuron variability than the OFF DBS condition (*p <* 0.001). This is evident in a visual comparison of the within-cycle SD of neuronal firing rates averaged across all cycles in the two conditions (Figure 6a), which shows a much higher mean SD on DBS. This coheres with our finding from the LFPCA model that level-1 (i.e., between-neuron) variability increases on DBS (Table 2). In evaluating the relationship between the variability and the mean FR across neurons within individual cycles (Figure 6c), we find that this increase in variability is associated with an increase in FR. That is, the increase in variability seen across cycles is likely due to increased neuronal firing in response to DBS. To understand whether this is attributable to changes at the individualneuron level, within-neuron variability must be explored.

**Fig. 6.**
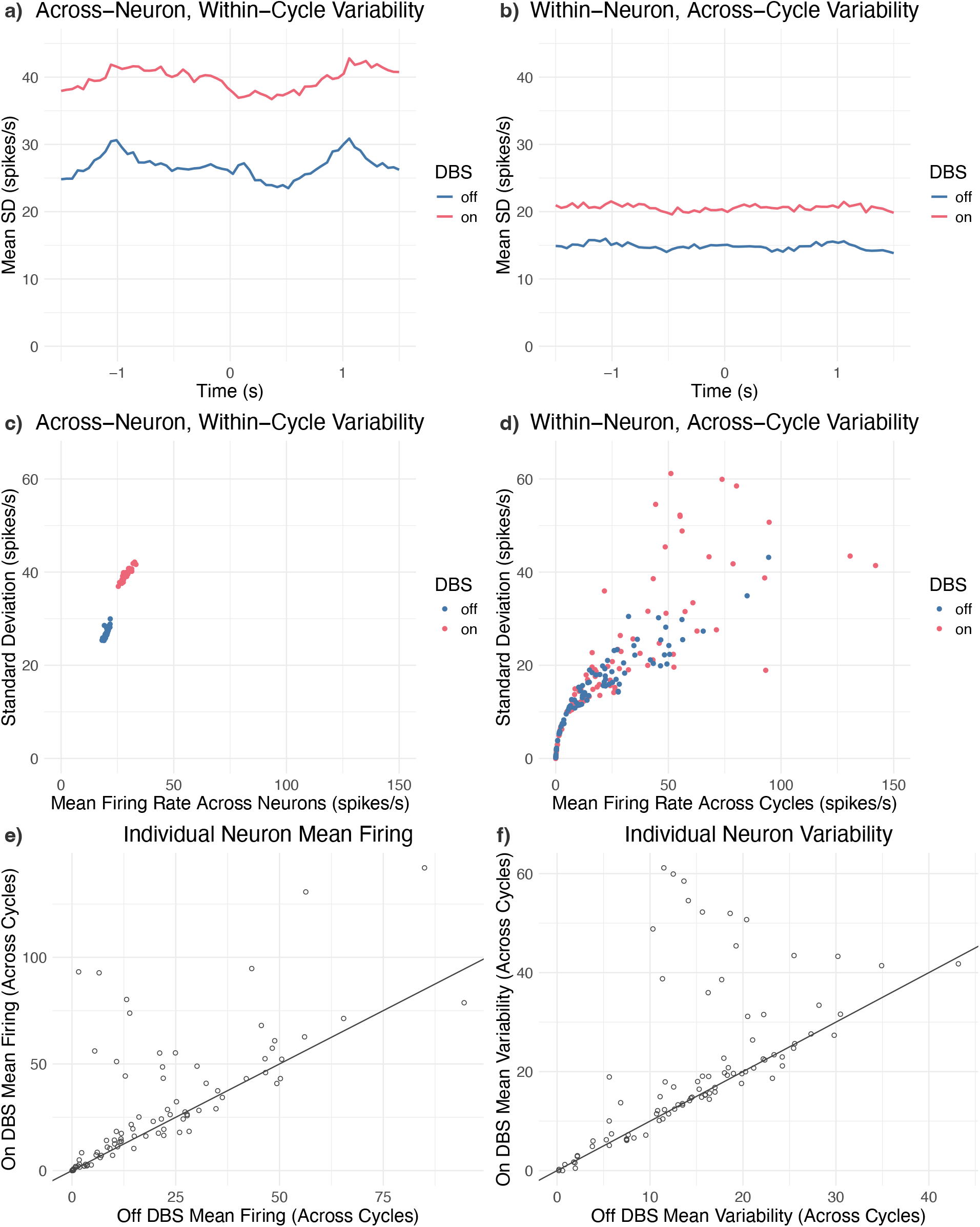
Figure comparing the variability of the firing rate OFF DBS (blue lines and dots) and ON DBS (pink lines and dots). Panel (a) displays the mean of the standard deviation of firing (spikes/s) across all neurons, within a given cycle, at each time point within a cycle. Panel (b) displays the mean of the standard deviation of firing (spikes/s) across all cycles, within a given neuron, at each time point within a cycle. Panel (c) displays the mean firing rate (spikes/s) and according standard deviation of the firing rate (spikes/s) across neurons for each individual cycle of data collection, both on and off DBS. Panel (d) displays the mean firing rate (spikes/s) and according standard deviation of the firing rate (spikes/s) across cycles for each individual neuron, both on and off DBS. Panel (e) displays the within-neuron mean firing rate across cycles (averaged over time) both OFF DBS (*x*-axis) and ON DBS (*y*-axis), such that each individual point represents an individual neuron’s mean firing rate in the two conditions. Panel (f) displays the within-neuron variability across cycles (averged over time) both OFF DBS (*x*-axis) and ON DBS (*y*-axis), such that each individual point represents an individual neuron’s average standard deviation in the two conditions.

In evaluating within-neuron variability, by calculating the standard deviation of the FR across all cycles within each neuron, a Wilcoxon signed rank test confirmed that the ON DBS condition had significantly higher across-cycle variability than OFF DBS (*p<* 0.001). This is evident through a visual comparison of the within-neuron SD of the firing rates averaged across all neurons in the two conditions (Figure 6b), which shows a much higher mean SD on DBS. To determine whether this is attributable to a specific subset of neurons, we assess the variability within individual neurons in Figure 6d (along the *y*-axis). It is evident that the increase in variability is largely *neuron-specific*. Specifically, there is a subset of neurons that have a notable increase in variability across cycles, such that their variability is far above the highest amount of variability seen in any of the neurons OFF DBS. This suggests that a number of neurons, but not all neurons, are changing non-trivially in their ON DBS firing activity. In contrast, another subset of neurons is maintaining a low across-cycle variability both ON and OFF DBS. Based on the information from the *x*-axis (mean firing rate) of Figure 6d, it is clear that many of these relatively stable neurons have a low or nearzero firing rate during DBS. The highly variable neurons, on the other hand, generally show a high rate of neuronal firing in response to DBS.

A comparison of the mean FR (Figure 6e) and the mean across-cycle variability (Figure 6f) at the neuron level further confirms that there are indeed a subset of neurons whose mean FR increases notably ON DBS relative to OFF DBS. Most neurons, however, display a similar rate of firing (either high or low) across the two conditions. A similar pattern is observed for mean variability, where a subset of neurons show a large increase in variability ON DBS relative to OFF DBS while most neurons show a relatively similar rate of variability across the two conditions. These neurons whose firing activity changes markedly on DBS would be of particular interest for further investigation in the future.

Overall, these single-neuron analysis results are consistent with our findings from the fast generalized LFPCA models that both level 1 and level 2 variability increase on DBS. From the LFPCA models, we found that level 2 variability increases notably on DBS and accounts for a much larger proportion of the total variance than in the OFF DBS condition, and the single-neuron results suggest that this increase is in fact driven by only a subset of neurons. While most neurons show relatively low variability across cycles and many also show a somewhat low mean activation off DBS, some neurons show a marked increase in across-cycle variability as well as a marked increase in mean FR during DBS (Figure 6d,e). This change in a subset of neurons also likely contributes to the increase in level 1 (between-neuron) variability that was also observed. In general, the high coherence between our findings of changes in variability from the LFPCA approach and those from a more “traditional” single-neuron spike analysis confirms that LFPCA can adequately uncover population-level patterns that result from changes in individual neuron behavior.

## Discussion

The current manuscript proposes a novel estimation technique for LFPCA, namely fast generalized LFPCA. This marginal approach leads to computational times that are orders of magnitude faster than the traditional technique proposed in Greven et al.(16), making the LFPCA model much more scalable for high-dimensional datasets. To exemplify its use, we apply the new fast generalized LFPCA approach to neuronal FR data to investigate population-level dynamics of neuronal activity. In applying LFPCA to FR data collected from a parkinsonian NHP both ON and OFF DBS, we aim to leverage the method to gain insight into the effect of DBS on population-level neuronal dynamics in a disease state, a relatively understudied topic in the field.

The fast generalized LFPCA technique proposed here has several adaptations that greatly improve computational efficiency. The primary gain comes from using a binning technique along the functional domain and fitting a marginal linear mixed model at each bin, which avoids the need for directly estimating and smoothing the covariance functions. Additionally, the variance components and score predictions are computed efficiently from closed-form estimators. Our simulation study results indicate that the proposed technique is significantly faster than traditional LFPCA, while having generally similar or better performance. We also note that as the number of observed time points increases, the computational efficiencies gained by the proposed fast technique become even more pronounced, making LFPCA scalable and widely applicable for a class of longitudinal functional data for which it was previously infeasible. Additionally, the generalized nature of the proposed technique makes it suitable for longitudinal functional data from any exponential family distribution.

In our application to neuronal FR data, we applied the fast generalized LFPCA approach to two separate data conditions: a block of FR data collected OFF DBS and a block of data collected ON DBS. In each condition, we extracted both between-neuron and within-neuron, across cycle eigenfunctions. Primarily, the LFPCA approach is useful as a dimension reduction tool to uncover large-scale patterns that exist in the data. The disentanglement of these two levels from LFPCA further allows to explore and understand the variation that exists both between neurons and across cycles. Here, we focus on the level-1, between-neuron eigenfunctions, which represent the broader population-level neuronal dynamics. We found that the vast majority (over 95%) of the total between-neuron variability could be explained by only a handful of eigenfunctions, greatly simplifying our interpretation of the shared patterns of the 90 unique neurons under study. In investigating the first four eigenfunctions, we found that the first component, explaining over 85% of the total between-neuron variability in both the ON and OFF DBS conditions, was a simple intercept-like component indicating whether a neuron was generally firing at a high versus low rate. Since the component score of each unique neuron in either condition indicates the overall FR of the neuron, scores can be compared across the two conditions to identify which neurons may be modulated by DBS (i.e., had a low rate of firing before DBS but a high FR on DBS, or vice versa). The subsequent eigenfunctions at the between-neuron level likely relate to the passive arm manipulation paradigm utilized during DBS administration, which was confirmed by relatively high correlations of components 2 and 3 with arm position (Section S4.2). Further investigation found that DBS does indeed modulate the relationship between arm position and neuronal FR. In particular, there was a change in both the overall relationship between arm position and firing, and in the specificity of the firing to particular components of motion (i.e., arm position versus arm velocity). These results show, as a proof of concept, how LFPCA can extract meaningful variation in the observed data and directly relate it to substantively meaningful information, such as pathophysiology of PD and the underlying mechanisms of DBS treatment.

Another benefit of the LFPCA approach is that it allows one to evaluate the overall variability in a condition block of data, and to separate out this variability into between-neuron (level 1) and within-neuron, across cycle (level 2) components, which are indistinct in a “standard” (single-level) PCA. We leveraged this ability across the OFF and ON DBS conditions to evaluate the effect of DBS on neuronal FR variability, and found that total variability increased on DBS. This was due to increased variability at both level 1 and level 2, but the relative increase at level 2 was particularly large. To determine whether this level 2 increase was due to an overall increased variability across cycles for all neurons or being driven by a specific subset of neurons, we conducted a post-hoc spike analysis of individual neuronal firing. This demonstrated that both across-neuron, within-cycle and and across-cycle, within-neuron variability increased significantly on DBS, and the increase in between-neuron variability was indeed driven by a particular subset of neurons. These neurons increased their overall mean firing, as well as the variability of their firing, in response to DBS. Other neurons, on the other hand, had relatively stable firing patterns both on and off DBS. One potential cause for this increase is antidromic activation, which past work has identified in approximately 30% of neurons in the M1 in response to STN DBS (the mechanism under study here) (24). Future work at the individual-neuron level should directly investigate how antidromic firing may relate to the increased activation and variability shown here, and LFPCA can assist in identifying particular neurons of interest. Our results demonstrate how LFPCA and single-neuron analyses can be used in conjunction to be mutually informative, while each provide unique information. Specifically, LFPCA is able to examine the temporal changes in coordinated activity within the neuronal population and their correlation with behavior, which cannot be established from single-neuron analyses alone.

There are several limitations of the present study that should be noted, each of which also lend themselves to a number of future directions. From a methodological perspective, our fast generalized LFPCA approach is not the only estimation technique that has been proposed to increase computational efficiency of LFPCA. Zipunnikov et al. (20) propose a method of moments approach for ultra high-dimensional data. However, their approach has limitations in modeling noise and larger sample sizes, and code implementation is not as readily available or familiar to `R`users. We also note that our simulation results showed that the proposed fast generalized LFPCA method had better recovery than traditional LFPCA, in terms of median ISE, for the level 1 (between-neuron) eigenfunctions and covariances, while the traditional approach had better recovery for level 2 (within-neuron) eigenfunctions and covariances. While the relative difference in performance was small, one might want to tune the bin selection in the fast approach more carefully if level 2 effects are primarily of interest.

From an applied perspective, we want to emphasize that while PCA-based methods are a popular approach for studying population-level neuronal dynamics, they are far from the only approach. GLMs (38, 39), spike word distributions (40, 41), and decoding approaches (42, 43) have all been used in the past to successfully uncover population-level patterns and provide their own potential insights into underlying dynamics. Decoding approaches in particular have been used in combination with arm movement information specifically, with the explicit aim of predicting external variables. The LFPCA approach does not directly relate the neuronal data to external information, and thus the decoding approach may be preferable if that is of primary interest. The LFPCA approach does, however, result in the low-dimensional representation of within-neuron, across-cycle variability in the form of the level 2 eigenfunctions, which contain more useful cycle-specific information. Here we did not directly explore the level 2 eigenfunctions, as our primary interest was in population-level (i.e., between-neuron) behavior. However, there are many ways the level 2 eigenfunctions could be used to explore within-neuron heterogeneity and differences that exist across cycles in future work focusing on individual-neuron behavior. Finally, we note that, due to inherent limitations in research based on NHPs, our sample size is only one subject, which limits generalizability. Although small sample sizes are relatively common when studying neuronal FR data, it will be important to replicate this approach in additional samples for making broader claims about mechanisms of DBS.

In summary, the present study provides a novel estimation technique for the LFPCA model, which greatly reduces computational time while maintaining accurate recovery of underlying effects. This technique is simple to implement for researchers familiar with any common mixed modeling software, and generalizable to any exponential family distribution, and thus may have utility in a wide range of substantive domains. Applying fast generalized LFPCA to neuronal FR data allows us to disentangle overall variability in neuronal FR into between-neuron variability and within-neuron, across-cycle variability, and highlights how this information can be used to understand population-level changes in neuronal activity. Our future research will further investigate how population-level dynamics relate to single-neuron analyses and the mutually informative and distinct information the two provide.

## Abbreviations

FR: firing rate
DBS: deep brain stimulation
MFPCA: multilevel functional principal component analysis
LFPCA: longitudinal functional principal component analysis

## ACKNOWLEDGEMENTS

The authors thank the Masonic Institute for the Developing Brain and the Department of Neurology, University of Minnesota, the National Institute of Health, and Minnesota’s Discovery, Research, and InnoVation Economy (MnDRIVE) Brain Conditions for their support of this research.

## Author contributions

**KAD:** Conceptualization, data curation, formal analysis, methodology, software, visualization, writing - original draft preparation, writing - review and editing. **NP:** Conceptualization, data curation, formal analysis, methodology, software, visualization, writing - original draft preparation, writing - review and editing. **DS:** Conceptualization, data curation, formal analysis, methodology, software. **GK:** Conceptualization, formal analysis, methodology, software, visualization, writing - original draft preparation, writing - review and editing. **AXO:** Project administration. **LZ:** Data curation, formal analysis, investigation, visualization. **JW:** Conceptualization, writing - review and editing, funding acquisition, supervision. **LAJ:** Writing - review and editing, funding acquisition, supervision. **JLV**: Writing - review and editing, funding acquisition, supervision. **MF:** Conceptualization, funding acquisition, methodology, supervision, writing - review and editing. **EC:** Conceptualization, methodology, software, funding acquisition, supervision, writing - review and editing.

## Supporting information

Additional supporting information may be found in the online version of the article at the publisher’s website. This consists of two components: a Supplementary Materials document (PDF) and a zip file of R code. The Supplementary Materials document contains additional results and details regarding analyses conducted in the Main Text. The zip file contains all code needed to fit the LFPCA model, perform the simulation study, and conduct all applied data analyses carried out in this manuscript.

