## Supplementary Materials for "Fast Generalized Longitudinal Functional Principal Components Analysis with Applications to Population-Level Neuronal Dynamics"

#### S1 Closed-form solutions for marginal linear-mixed models

First, we obtain estimates of the variance components. We collapse the model to a cell means model to reduce the number of steps. Define $\bar{R}^b_{ij} = \frac{1}{L_b}\sum_{l=1}^{L_b} \hat{R}^b_{ijl}$, i.e., the means within each bin for each subject-visit. The collapsed model is $\hat{R}^b_{ij} = \boldsymbol{Z}_{ij}^\top \boldsymbol{B}^b_i + u^b_{ij}$, where $u^b_{ijl} = W^b_{ij} + \frac{1}{L_b}\sum_{l=1}^{L_b} \epsilon^b_{ijl}$; in vector form, $\bar{\boldsymbol{R}}^b_i = \boldsymbol{Z}_i \boldsymbol{B}^b_i + \boldsymbol{u}^b_i$. We obtain an estimate of the error variance as

$$\hat{\sigma}^2_b = \frac{\sum_{i=1}^I \sum_{j=1}^J \sum_{l=1}^{L_b} (\hat{R}^b_{ijl} - \bar{\hat{R}}^b_{ij})^2}{IJ(L_b - 1)}.$$

Let $\omega^2_b$ be the variance of $u^b_{ijl}$, so $\omega^2_b = \tau^2_b + \frac{\sigma^2_b}{L_b}$. We estimate $\omega^2_b$ using a standard OLS estimator on the collapsed model, that is,

$$\hat{\omega}^2_b = \sum_{i=1}^I \frac{||\bar{\boldsymbol{R}}^b_i - \boldsymbol{Z}_i \hat{\boldsymbol{B}}^b_i||^2}{I(J-2)},$$

where $\hat{\boldsymbol{B}}^b_i = (\boldsymbol{Z}_i^\top \boldsymbol{Z}_i)^{-1} \boldsymbol{Z}_i^\top \bar{\boldsymbol{R}}^b_i$. Then, we estimate $\tau^2$ as $\hat{\tau}^2 = \hat{\omega}^2_b - \frac{\hat{\sigma}^2_b}{L_b}$. Finally, $\Sigma_b$ is estimated by

$$\hat{\Sigma}^b = \frac{1}{I}\sum_{i=1}^I \hat{\boldsymbol{B}}^b_i (\hat{\boldsymbol{B}}^b_i)^\top - \hat{\omega}^2_b (\boldsymbol{Z}_i^\top \boldsymbol{Z}_i)^{-1}.$$

After the variance components are estimated, we obtain predictions of $\boldsymbol{B}^b_i$ and $W^b_{ij}$ via Henderson's mixed model equations (MME).

#### S2 MFPCA

##### S2.1 Multilevel Functional PCA (mfPCA) and Fast mfPCA

We first briefly outline the technical and mathematical details of the mfPCA approach. For a full, in-depth exploration of mfPCA, see Di et al. (2009) and Chapter 8 of Crainiceanu et al. (2024). For a full treatment of the recent advancement of "fast mfPCA," see Cui et al. (2023). Consider multilevel functional data of the form $R_{ij}(t)$ for neuron $i = 1, \ldots, I$ and cycle $j = 1, \ldots, J_i$ at time $t \in \{t_1, \ldots, t_L\} \in \mathcal{T}$, where $\mathcal{T}$ is a compact domain. Note that each function has $L$ observations at the same set of time points. We can then consider a functional ANOVA model, with measurement error, for this data of the form $R_{ij}(t) = S_{ij}(t) + \epsilon_{ij}(t) = \mu(t) + \eta_j(t) + A_i(t) + B_{ij}(t) + \epsilon_{ij}(t)$, where $\mu(t)$ is the population mean function, $\eta_j(t)$ is the $j$th cycle-specific shift from $\mu(t)$, $A_i(t)$ is the neuron-specific mean deviation for the $i$th neuron under study, $B_{ij}(t)$ is the random $j$th cycle deviation for the $i$th neuron's mean $A_i(t)$, and $\epsilon_{ij}(t)$ is random white-noise error with a variance of $\sigma^2$. Note that the random functions $A_i(t)$ and $B_{ij}(t)$ are assumed to be mutually independent processes with a mean of 0 and covariance functions of $K_B(t, u) = \text{cov}\{A_i(t), A_i(u)\}$ and $K_W(t, u) = \text{cov}\{B_{ij}(t), B_{ij}(u)\}$, which give the between-neuron and within-neuron variance, respectively. We can then define the total variance of the smooth functional data as $K_T(t, u) := \text{cov}\{S_{ij}(t), S_{ij}(u)\} = K_B(t, u) + K_W(t, u)$.

Following Mercer's theorem (Mercer, 1909), the between-neuron covariance, $K_B$, can be eigendecomposed as $K_B(t, u) = \sum_{k \geq 1} \lambda_k^{(1)} \phi_k(t)\phi_k(u)$, where $\lambda_1^{(1)} \geq \lambda_2^{(1)} \geq \cdots \geq 0$ are the (level-1) eigenvalues that are associated with according (level-1) orthonormal eigenfunctions $\phi_k(t)$, such that $\int_{\mathcal{T}} \phi_{k_1}(t)\phi_{k_2}(t)dt = \delta_{\{k_1 = k_2\}}$ for any pair $(k_1, k_2)$, where $\delta_{\{\cdot\}}$ is an indicator function (i.e., is equal to 1 if $k_1 = k_2$ and 0 otherwise). Following the Karhunen-Loeve theorem (Karhunen, 1947), the neuron-specific random functions $A_i(t)$ can then be written as $A_i(t) = \sum_{k \geq 1} \xi_{ik}\phi_k(t)$, where $\xi_{ik}$ are mutually uncorrelated scores with a mean of zero and a variance of $\lambda_k^{(1)}$. The within-neuron (across cycle) variance can be eigendecomposed in a similar fashion as $K_W(t, u) = \sum_{k \geq 1} \lambda_k^{(2)} \psi_k(t)\psi_k(u)$, where $\lambda_1^{(2)} \geq \lambda_2^{(2)} \geq \cdots \geq 0$ are the (level-2) eigenvalues that have associated orthonormal (level-2) eigenfunctions $\psi_k(t)$. The cycle-and-neuron-specific random functions $B_{ij}(t)$ can then be written as $B_{ij}(t) = \sum_{k \geq 1} \zeta_{ijk}\psi_k(t)$, where $\zeta_{ijk}$ are mutually uncorrelated scores with a mean of zero and a variance of $\lambda_k^{(2)}$. To achieve dimensionality reduction, the first several principal components are kept at each level, where the number can be chosen, for example, based on proportion of variance explained (PVE) (Di et al., 2009; Cui et al., 2023). Through such decomposition, mfPCA reduces the multilevel functional data into two sets of uncorrelated scores: the level-1 (between-neuron) scores $\xi_{ik_1}$ and the level-2 (within-neuron, between-cycle) scores $\zeta_{ijk_2}$. We also obtain eigenfunctions at level-1 and level-2, which show the patterns of variation in the functional data at each level. Overall, the mfPCA approach provides an explicit decomposition of the between-neuron variability and the within-neuron, between-cycle variability.

The mfPCA approach is well-suited to model the repeated measures from multiple neurons in our motivating SF data, as it allows the explicit investigation of population-level neural activity (by focusing on the level-1 eigenfunctions) while appropriately accounting for the heterogeneity between individual neurons and across cycles within neurons. However, the application of the approach has been limited in the past due to computational constraints. Specifically, the traditional mfPCA model requires a number of computations that is proportional to the *cube* of the number of observations $L$. As $L$ increases, computation time thus increases substantially. However, recent developments (Cui et al., 2023) enable a substantially faster model fitting approach, referred to as "fast mfPCA." The estimation approach for fast mfPCA is outlined in Algorithm S1. Compared to mfPCA, fast mfPCA makes computational gains through several different areas. First, by transforming functional data such that the underlying smooth curves have the desired covariance operators, it substantially reduces the number of computations needed to construct smooth covariance matrices compared to the traditional method of moments approach. Indeed, using the fast covariance estimation (FACE (Xiao et al., 2016)) method on the transformed data to estimate the covariance operators avoids direct eigenanalysis of the covariance matrices, requiring only a number of computations that is proportional to $L$ rather than $L^3$. In addition, rather than using the best linear unbiased prediction (BLUP), the principal component scores are predicted through mixed model equations (MME) (Henderson, 1973) in fast mfPCA, which are computationally more efficient. Ultimately, fast mfPCA requires only a number of computations that scales *linearly* with $L$, which is two orders of magnitude faster than the traditional mfPCA approach.

The computational efficiencies of fast mfPCA make it capable of modeling data that is much more high-dimensional than what the traditional mfPCA can handle. Combined with its easily accessible software implementation, fast mfPCA has substantially broadened the range of functional data applications, including our analysis of high-dimensional SF data. Here, we illustrate how this approach can be used to draw conclusions about population-level functional dynamics and treatment paradigms (e.g., DBS), while noting that mfPCA is broadly applicable to many other data types and substantive questions.

**Algorithm S1:** Fast MFPCA Algorithm (Cui et al., 2023)

1: Estimate the mean functions $\mu(t)$ and $\eta_j(t)$ and subtract them from the observed data
2: Apply FACE to the functional data, transformed as $\sqrt{nw_i}(\tilde{\mathbf{R}}_{ij}), 1 \leq j \leq J_i, 1 \leq i \leq I)$, for total covariance
3: Apply FACE to the functional data, transformed as $\sqrt{nv_iJ_i}(\tilde{\mathbf{R}}_{ij} - \bar{\mathbf{R}}_i), i \leq j \leq J_i, 1 \leq i \leq I$, for within-neuron covariance and obtain within-neuron eigenfunctions and eigenvalues
4: Use the difference between total and within-neuron covariance from above steps to calculate between-neuron covariance and extract between-neuron eigenfunctions and eigenvalues
5: Estimate error variance $\sigma^2$ by $\hat{\sigma}^2 = \int_{\mathcal{T}}\{\hat{K}_S(t,t) - \bar{K}_S(t,t)\}dt$
6: Estimate scores using Henderson's mixed model equations (see Eq. 4 of Cui et al. (2023))

### S3 Additional Results: Simulation Study

#### S3.1 Covariance Functions Recovery

Figure S1 presents the the recovery (in terms of integrated squared error [ISE]) of the covariance functions in the simulation study presented in the Main Text (Section 4).

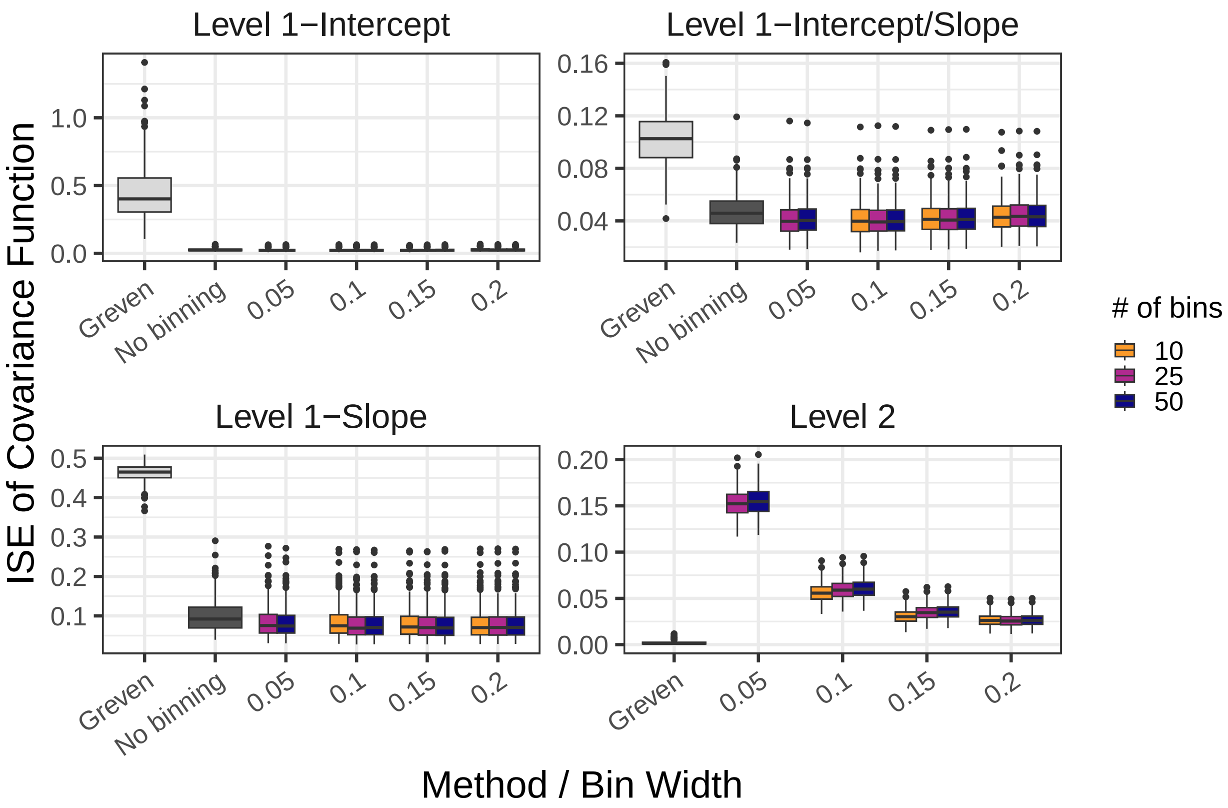


Figure S1: Boxplots of integrated squared error (ISE) of the covariance functions in simulation.

### S4 Additional Results: Application

#### S4.1 Level-2 LFPCA Eigenfunctions

We present here the first four eigenfunctions that explain the greatest amount of variance at level-2 in both the OFF DBS condition (Figure S2) and the ON DBS condition (Figure S3. Recall that these level-2 eigenfunctions are at the within-neuron, between-cycle level. Note that the $y$-axis across Figures S2 and S3 is scaled to be consistent across the conditions.

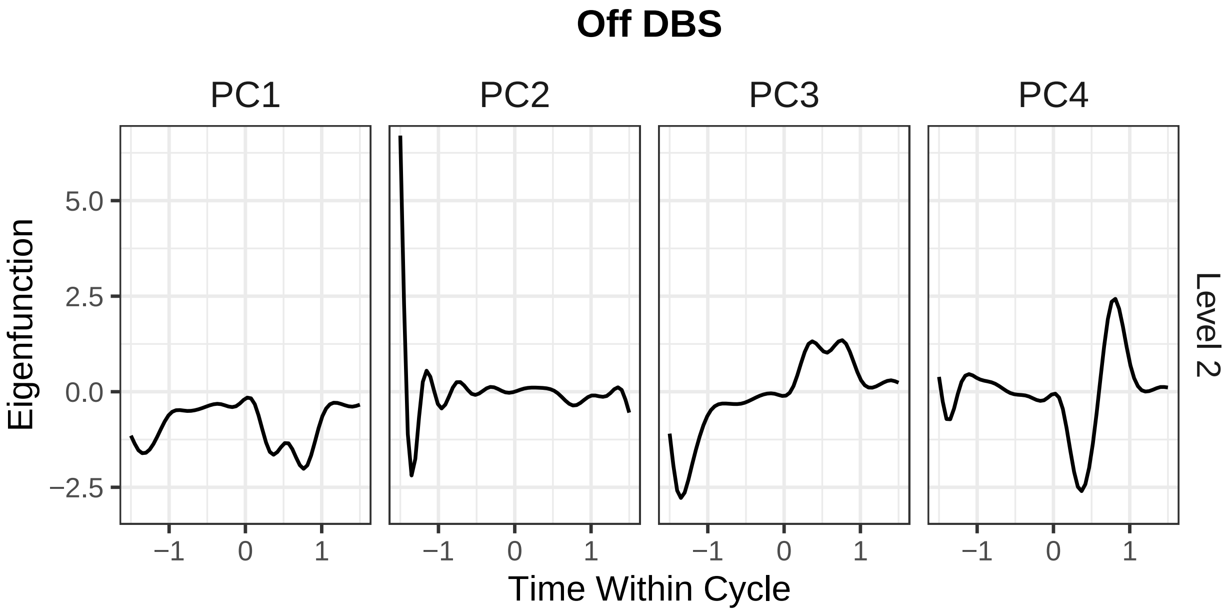


Figure S2: The first four eigenfunctions (PCs) explaining the largest proportion of variance at level-2 of the OFF DBS condition of activity. The figures display the first four eigenfunctions of the within-neuron, across cycle intercepts.

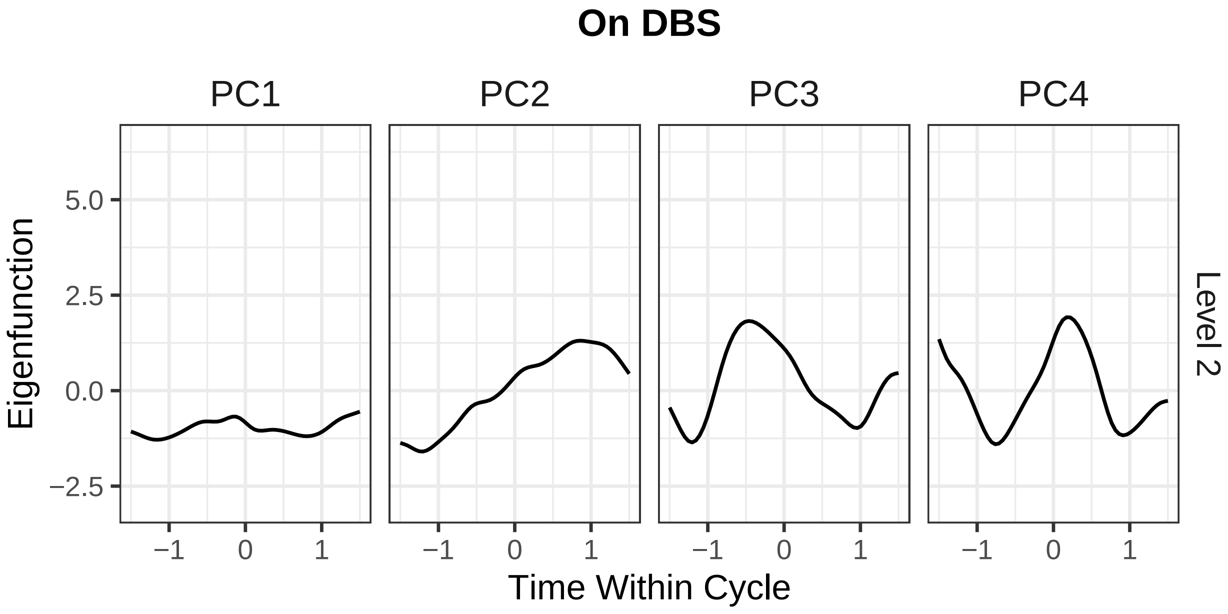


Figure S3: The first four eigenfunctions (PCs) explaining the largest proportion of variance at level-2 of the ON DBS condition of activity. The figures display the first four eigenfunctions of the within-neuron, across cycle intercepts.

#### S4.2 Correlations With Passive Arm Movement

##### S4.2.1 Methods: Correlations of Eigenfunctions with Passive Arm Manipulation

To further interpret and better understand the estimated eigenfunctions, we explore their relationship with the kinematic data from the simultaneous passive arm manipulation. Because we are primarily interested in understanding population-level neural dynamics, we only explore the relationship with the level-1 (i.e., between-neuron) eigenfunctions. However, we note that the level-2 eigenfunctions could be further investigated and interpreted as well, particularly if the focus is on between-cycle or within-neuron effects.

Let the arm position at each discretized time point of data collection $t$, for a given cycle $j$, be denoted as $x_j(t)$ for the arm position in the X (Anterior-Posterior) direction, $y_j(t)$ for the position in the Y (Superior-Inferior) direction, and $z_j(t)$ for the position in the Z (Lateral-Medial) direction.

From the initial position values $x_j(t)$, $y_j(t)$, and $z_j(t)$, we can additionally calculate velocity in the same X, Y, and Z directions by taking the derivative of the specific position with respect to time. These velocity values are denoted as $x'_j(t)$, $y'_j(t)$, and $z'_j(t)$.

We will let the total collection of kinematic measures of interest (arm position and arm velocity in each direction) be denoted as $M_j = \{x_j(t), y_j(t), z_j(t), x'_j(t), y'_j(t), z'_j(t)\}$. For each measure $m_{jh} \in M_j, h = 1, .., 6$, we then calculate the average value across the $J$ cycles to obtain a single value at each time point. That is, using the arm position in the X direction as an example, we calculate $\bar{x}(t) = \sum_{j=1}^{J} x_j(t)/J$ to obtain the average arm position in the $X$ direction across cycles. We then repeat this for each outcome, such that there is a single function across the $t$ time points for each outcome of interest, to obtain $\bar{M} = \{\bar{x}(t), \bar{y}(t), \bar{z}(t), \bar{x}'(t), \bar{y}'(t), \bar{z}'(t)\}$. We then calculate the Pearson sample correlation (denoted by $r$) of the first 3 level-1 eigenfunctions with each measure $\bar{m}_h(t) \in \bar{M}(t)$, to obtain $r_{\bar{m}_h(t),\phi_1(t)}, r_{\bar{m}_h(t),\phi_2(t)}$, and $r_{\bar{m}_h(t),\phi_3(t)}$, where $\phi_1(t)$ is the level-1 eigenfunction explaining the greatest proportion of variance, $\phi_2(t)$ is the eigenfunction explaining the second greatest proportion of variance, and $\phi_3(t)$ explains the third greatest proportion of variance.

##### S4.2.2 Methods: Permutation Test for Between-Condition Differences

To evaluate whether the kinematic correlations differ across the DBS conditions, we implement a permutation test. Where $\rho$ denotes Pearson correlation of the population, we specify our null hypothesis as no difference between the two conditions and our alternative hypothesis as non-zero difference (that is, $H_0 : |\rho^{OFF}_{\bar{m}_h(t),\phi_k(t)}| - |\rho^{ON}_{\bar{m}_h(t),\phi_k(t)}| = 0$ and $H_A : |\rho^{OFF}_{\bar{m}_h(t),\phi_k(t)}| - |\rho^{ON}_{\bar{m}_h(t),\phi_k(t)}| \neq 0$ for $h = 1, \ldots, 6,\ k = 1, 2, 3$). We use $|\rho^{OFF}_{\bar{m}_h(t),\phi_k(t)}| - |\rho^{ON}_{\bar{m}_h(t),\phi_k(t)}|$ rather than $\rho^{OFF}_{\bar{m}_h(t),\phi_k(t)} - \rho^{ON}_{\bar{m}_h(t),\phi_k(t)}$ for our null hypothesis due to the sign indeterminacy of eigenfunctions. The functional nature of the data makes derivation of the theoretical/asymptotic null distribution for the $|\rho^{OFF}_{\bar{m}_h(t),\phi_k(t)}| - |\rho^{ON}_{\bar{m}_h(t),\phi_k(t)}|$ difficult, but tests of the distribution can instead be carried out using nonparametric approaches (Ramsay and Silverman, 1997; Ferraty and Vieu, 2006; Crainiceanu et al., 2012), such as permutation tests (Corain et al., 2014). Specifically, we can implement a permutation test that pairs $\phi_k$ (see Section 5.1) with $\bar{m}_h(t)$ for each combination of $k$, $h$ (see Section S4.2.1). Our permutation procedure for the test is described in Algorithm S2.

Step 0 is to compute the observed test statistic. The correlation between the eigenfunction of interest and the averaged position trajectory is calculated for each condition, as described in Section 2.3. Then, absolute values of correlations and their difference are computed. Steps 1-2 are then to generate the null distribution of test statistic. On each permutation iteration, $(j, \text{DBS})$ pairs are randomly shuffled for SF and movement data together, maintaining the original number of cycles per condition and preserving the relationship between SF activity and motion within cycles. Note that as neurons within each $(j, \text{DBS})$ pair are shuffled together, the neuron-level correlation structure within each cycle is preserved, and this permutation strategy implicitly assumes independence of cycles. Then, the same correlation computations used on the original blocks are performed on the permuted blocks to obtain the difference in correlation between the permuted ON and OFF DBS blocks.

In our analysis, Steps 1-2 of the permutation algorithm were repeated 1,000 times (i.e., $B = 1000$) to obtain null distributions of $D_{hk}$ under the assumption that condition assignment does not affect the correlation between each position trajectory and eigenfunction. See Figure S4 for histograms of these null distributions. From these null distributions, we compute permutation-based $p$-values, and correct them for multiple comparisons using a False Discovery Rate (FDR) correction (Benjamini and Hochberg, 1995). We perform the multiple comparison correction separately for each eigenfunction, adjusting $p$-values across the six kinematic measures associated with that eigenfunction.

###### S4.2.3 Results: DBS Modulates the Relationship Between Eigenfunctions and Passive Movement

In addition to exploring and understanding the variability between neurons and across cycles, and the effect of DBS on those outcomes, we can also leverage the LFPCA output to better understand broad, population-level patterns in neuronal firing. Specifically, to better understand how the level-1 eigenfunctions can inform our understanding of large-scale neuronal firing patterns, we explore their relationship with external information; namely, the arm position information collected as part of the passive manipulation protocol described in Section **??**. As the neuronal spike firing information and arm motion information were collected simultaneously, their contemporaneous relationship can be explored. We first calculate the Pearson correlation between the first three level-1 eigenfunctions and the arm position values (averaged across cycles) at each time point across the data collection cycle (Table S1). The arm position was recorded in the X (Anterior-Posterior), Y (Superior-

**Algorithm S2:** Permutation Algorithm

**Input:** SF data $\{R_j^{OFF},\ j=1,\dots,J^{OFF}\}$ and $\{R_j^{ON},\ j=1,\dots,J^{ON}\}$ where $R_j=\{R_{ij},\ i=1,\dots,I\}$; arm movement data $\{M_j^{OFF},\ j=1,\dots,J^{OFF}\}$ and $\{M_j^{ON},\ j=1,\dots,J^{ON}\}$; number of permutations $B$

**Output:** $p$-values for the hypothesis tests $H_0: |\rho^{OFF}_{\bar{m}_h(t),\phi_k(t)}|-|\rho^{ON}_{\bar{m}_h(t),\phi_k(t)}|=0$ with $H_A: |\rho^{OFF}_{\bar{m}_h(t),\phi_k(t)}|-|\rho^{ON}_{\bar{m}_h(t),\phi_k(t)}|\neq 0$ for $h=1,\dots,8,\ k=1,2,3$

0: Compute observed test statistic

(a) Fit fast generalized LFPCA on $\{R_j^{OFF},\ j=1,\dots,J^{OFF}\}$ and $\{R_j^{ON},\ j=1,\dots,J^{ON}\}$ to obtain $\phi_k^{OFF}(t),\ \phi_k^{ON}(t),\ k=1,2,3$ respectively (Section 5.1)

(b) Obtain $\bar{M}^{OFF}$ and $\bar{M}^{ON}$ (Section **??**)

(c) Compute $D_{hk}=|r^{OFF}_{\bar{m}_h(t),\phi_k(t)}|-|r^{ON}_{\bar{m}_h(t),\phi_k(t)}|$ for $h=1,\dots,8,\ k=1,2,3$ (Section **??**)

**For** $b=1,\dots,B$**, do:**

1: Pool SF and movement data across both DBS conditions. For each pair $(R_j^{DBS}, M_j^{DBS})$, randomly shuffle $(j,\mathrm{DBS})$ pairs (where "DBS" is either "ON" or "OFF" label) across the $J^{OFF}+J^{ON}$ total cycles. Separate permuted data by DBS label, forming $\{R_{j'}^{OFF},\ j'=1,\dots,J^{OFF}\}$, $\{M_{j'}^{OFF},\ j'=1,\dots,J^{OFF}\}$, $\{R_{g'}^{ON},\ g'=1,\dots,J^{ON}\}$, $\{M_{g'}^{ON},\ g'=1,\dots,J^{ON}\}$

2: Repeat steps 0a–0c with permuted data from step 1 to obtain permuted test statistic $D'_{hk}$

**End**

3: Compute the $p$-value as the proportion of the permuted test statistics that were as extreme as or more extreme than the observed test statistic

Inferior), and Z (Lateral-Medial) directions, and velocity was also calculated in these same directions.

In examining the correlation values, we see that the first component ("E1") shows strong correlations with the velocity values (particularly in the Z direction) and little-to-no correlation with the position values in the OFF DBS condition. During DBS, the correlations with the position values get notably stronger, while the correlations with the velocity values get weaker. We note that as many of the correlations indicate a statistically significant relationship between kinematics and the eigenfunctions, we focus here primarily on the relative magnitudes of the correlations. The second component ("E2") shows moderate-to-strong correlations with the position information and strong correlations with the velocity values in the OFF DBS condition. During DBS, the correlations with the position and velocity values drop significantly, becoming null-to-moderate in all directions. For the third component ("E3"), we see a different pattern: strong correlations with the position information (especially in the Z direction) in the OFF DBS condition and moderate-to-strong correlations with the velocity values. During DBS, the correlations with the position values get weaker, while the correlations with the velocity values get stronger.

To determine whether the observed differences in correlations across the OFF DBS and ON DBS conditions were significant, we implemented a permutation test, as described in Section S4.2.2. Following an FDR correction for multiple comparisons, significant differences were detected for the correlation between the second component ("E2") and the Z (Lateral-Medial) position and velocity, as well as the Y (Superior-Inferior)

Table S1: Correlation between the first three eigenfunctions explaining the greatest proportion of variance ("E1", "E2", and "E3") and the wrist position information (in the X [Anterior-Posterior], Y [Superior-Inferior], and Z [Lateral-Medial] directions) and velocity information (in the same X, Y, and Z directions). The correlation for the first three eigenfunctions are reported for the separately-constructed components from the OFF deep brain stimulation (DBS) and ON DBS conditions of activity. Symbols indicate false discovery rate (FDR-corrected) significance values of the test of the Pearson correlation with each kinematic value and the eigenfunction in that condition. The highlighted rows represent the (absolute) difference in the correlations across the OFF DBS and ON DBS conditions. Symbols indicate FDR-corrected significance values based on our permutation test for evaluating the significance of the difference in the correlation values across conditions.

| | **X (Ant.-Post.)** | **Y (Sup.-Inf.)** | **Z (Lat.-Med.)** | **X Velocity** | **Y Velocity** | **Z Velocity** |
|---|---|---|---|---|---|---|
| **E1** | | | | | | |
| OFF DBS | -0.127 | -0.252 | 0.058 | 0.864*** | 0.616*** | -0.936*** |
| ON DBS | -0.488*** | -0.343* | 0.607*** | -0.791*** | -0.836*** | 0.644*** |
| Difference | 0.361 | 0.091 | 0.549 | 1.655 | 1.452 | 1.580 |
| **E2** | | | | | | |
| OFF DBS | 0.529*** | 0.615*** | -0.463*** | -0.775*** | -0.638*** | 0.801*** |
| ON DBS | -0.106 | -0.110 | 0.148 | 0.054 | -0.022 | -0.140 |
| Difference | 0.635 | 0.725 | 0.611* | 0.829 | 0.615* | 0.941* |
| **E3** | | | | | | |
| OFF DBS | -0.825*** | -0.740*** | 0.854*** | -0.600*** | -0.669*** | 0.510*** |
| ON DBS | -0.656*** | -0.508*** | 0.757*** | -0.707*** | -0.792*** | 0.550*** |
| Difference | 0.169 | 0.233 | 0.097 | 0.107 | 0.123 | 0.040 |

* = $p < 0.05$, ** = $p < 0.01$, *** = $p < 0.001$.

velocity. Looking at Table S1, we can see that all of these are due to decreased magnitudes in correlations from OFF DBS to ON DBS. That is, for E2, the correlation with the Z position and Z and Y velocities significantly declined during DBS. This change was strongest for the correlation between E2 and the Z arm velocity, where the overall relationship changes from highly significant and positive OFF DBS to non-significant ON DBS. There were, though, components and directions that showed increased in correlations ON DBS, such as E3 and velocity in all directions. Interestingly, the increase in correlation does seem to be specific to velocity for E3, as the correlations with position decrease in all directions ON DBS. However, these did not reach a threshold of statistical significance ($p$s $< 0.07$ for Y and Z velocity and X and Z position).

We note that E1 did not show any significant change in relation to the kinematic data during DBS. E1 did show relatively high correlations with some kinematic values, which is somewhat surprising given its apparently flat shape. We attribute this primarily to the slight sinusoidal shape of the function, which although very small in magnitude, may be sufficient to induce correlations with the kinematic variables. This may also have led to the reversal in the direction of the correlation that we see between blocks, which leads to what appear to be very large changes in magnitude between the two blocks. However, the permutation test did not find this differences to be significant, suggesting that despite the apparent large magnitude, E1 is not changing notably between blocks and is fairly consistent across the time domain. This supports the interpretation of this component as primarily an indicator of general neuronal firing.

Overall, we see that DBS is significantly modulating the relationship between neuronal firing and passive arm movement at the between-neuron level. For E2, this has the effect of decreasing the correlation between neuronal firing and arm position and velocity. For E3, this has the effect of increasing the specificity of the kinematic correlations, such that correlations with arm position are stronger off DBS and correlations with arm velocity are stronger on DBS.

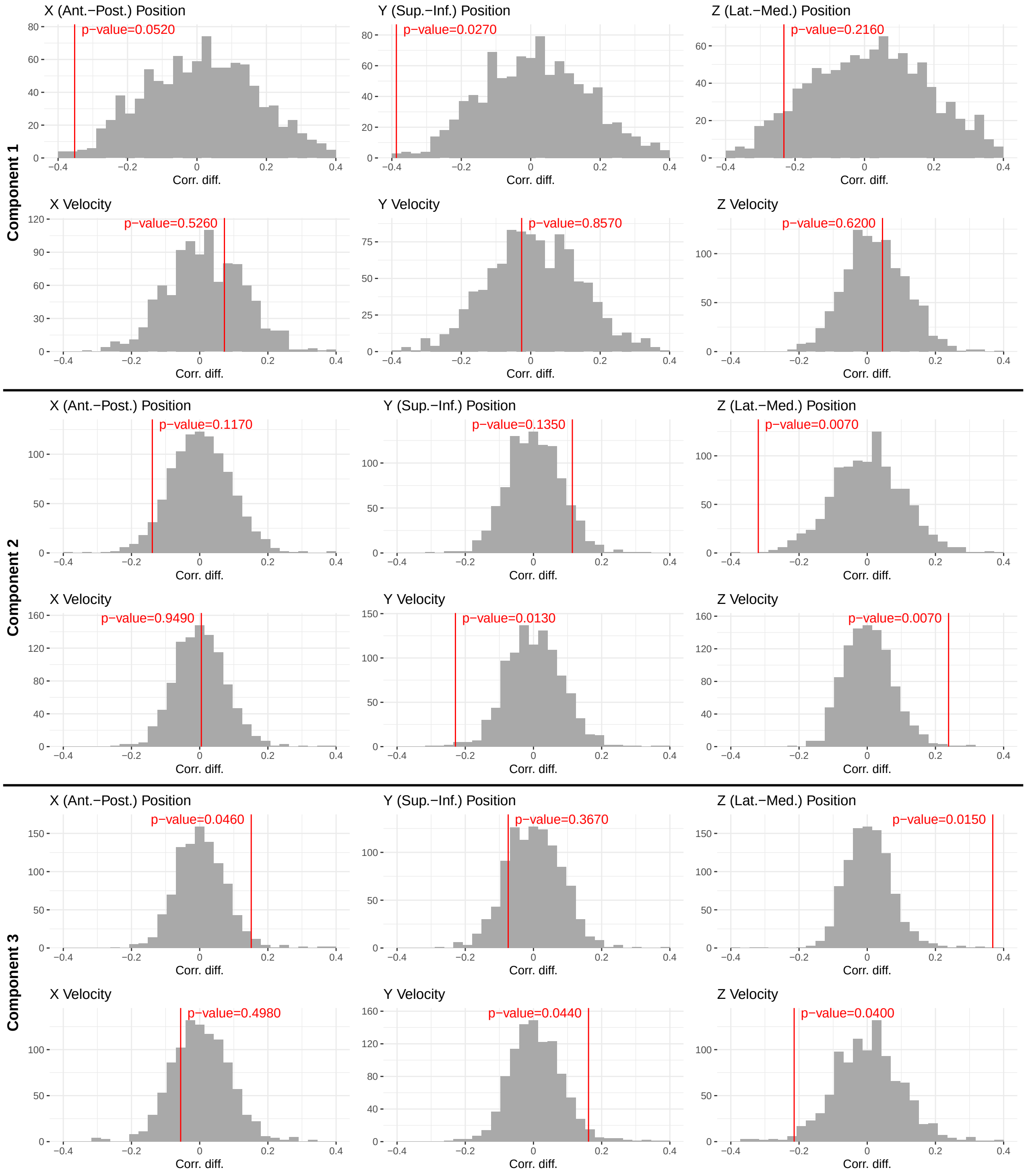


Figure S4: Permutation-based histograms of null distributions of the absolute difference in Pearson correlations between the first three level-1 eigenfunctions obtained with the fast generalized LFPCA model and arm movement measures, comparing the OFF DBS and ON DBS conditions. The figure is split into three horizontal sections for Eigenfunctions 1–3 ("Component 1", "Component 2", and "Component 3", respectively); each section contains eight histograms with panel titles indicating the measure (X, Y, Z position; X, Y, Z velocity). In each histogram, the $x$-axis is the absolute difference in correlation and the $y$-axis is count; the red vertical line marks the observed value and the annotation gives the permutation-based (uncorrected) p-value, which was computed as the proportion of the permuted correlation differences that were as extreme as or more extreme than the observed difference.
